# Complete genomes reveal the full extent of *Mycobacterium tuberculosis* complex diversity across evolutionary scales

**DOI:** 10.1101/2025.08.05.667821

**Authors:** Ana María García-Marín, Manuela Torres-Puente, Llúcia Martínez-Priego, Griselda De Marco, Miguel Moreno-Molina, Martin Hunt, Zamin Iqbal, Ana Gil-Brusola, Valencia Region TB Working Group, Mariana G. López, Fernando González-Candelas, Javier Alonso-del-Real, Iñaki Comas

## Abstract

Advances in short-read sequencing have enhanced our understanding of *Mycobacterium tuberculosis complex* (MTBC), but fail to capture its complete genomic diversity. We applied long-read sequencing to 216 isolates from the Valencia Region (Spain) and generated high-quality, complete genomes, revealing detailed insights into MTBC evolution across timescales. Complete genome comparisons increased the estimated evolutionary rate by 1.5-fold, resulting in a median of 312 (–1 to 792) additional SNPs per pairwise comparison. Multiple diversity hotspots were identified, mostly in the *pe/ppe* genes and driven by gene conversion. However, most PE/PPE epitopes were hyperconserved, with notable exceptions involving vaccine candidates. Incorporating previously undetected SNPs and indels improved resolution in transmission analyses. Furthermore, patient-specific reference mapping validates only 5–10% of within-host diversity detected by standard pipelines, indicating substantial overestimation in previous studies. These findings expand our view of MTBC diversity and have important implications for understanding host-pathogen interactions, epidemiology, and transmission dynamics.

## INTRODUCTION

Despite advances in diagnosis, surveillance, and treatments, tuberculosis (TB) remains a major global health problem^1^. The widespread use of short-read sequencing has enabled the rapid and large-scale sequencing of the *Mycobacterium tuberculosis* complex (MTBC), providing new insights into its genetic diversity. This has fueled extensive efforts to link MTBC genetic variation to key phenotypes such as virulence, transmission, drug resistance, host tropism, and disease manifestations^2^.

However, genomic studies are hampered by the limitations of short-read sequencing in resolving repetitive or highly complex regions and structural variants, which are excluded from bioinformatic analyses due to mapping challenges^3^. As a consequence, approximately 5-10% of the genome cannot be accurately assessed and remains understudied^4,5^. Paradoxically, some of the most diverse and functionally relevant regions of the genome, including *pe/ppe* genes, are inaccessible by short-read sequencing.^5–7^ Those regions also harbor the potential to increase the resolution of transmission and within-host variation studies. We are therefore blinded to regions that can potentially improve our understanding of tuberculosis disease and epidemiology.

Long-read sequencing technologies are currently the only viable alternative to overcome these limitations since they allow for the accurate reconstruction of the complete MTBC genome through *de novo* assembly^3^. Previous studies using MTBC assemblies were constrained by small sample sizes and the moderate quality of long-read data, often requiring polishing with short reads or hybrid assembly approaches^8–10^. Still, progress has been made to address specific questions, although sample sizes preclude the generalization of results in many cases^11–13^. Finally, while the accuracy of long-read sequencing has improved over the last few years^14^, sequencing results are still limited by the low quality of bacterial DNA (resulting from harsh conditions of extraction protocols) and the high cost of these technologies^15^.

Here, we present the largest dataset of high-quality complete MTBC genomes published to date, comprising 216 MTBC clinical isolates collected in 2016 from the Valencia Region, Spain, with matched short-read and genome assembly data. We generated complete genome sequences solely from long-read HiFi sequencing data, achieving exceptional quality without including short reads for polishing or performing hybrid approaches.

Compared to previous analyses, our results reveal substantial additional genetic diversity across the MTBC, with a particular focus on Lineage 4. Most of this diversity is associated with the PE/PPE gene family, a largely uncharacterized group of genes in MTBC. Point mutations and structural variations contribute to that diversity, but it is mainly driven by gene conversion events, contrary to what is observed in the rest of the genome. However, we show that the epitopes in PE/PPE antigens are hyperconserved, with some salient exceptions. Furthermore, our results provide new insights to redefine transmission boundaries based on genetic distances. Finally, the use of a complete genome from the same patient as a reference reveals a substantial reduction in non-fixed variation when mapping short reads from serial isolates, prompting a reassessment of current within-host diversity estimates and the dynamics of *Mycobacterium tuberculosis* during infection.

## METHODS

### Study design

We performed long-read sequencing on all MTBC culture-positive samples collected as part of a local population-based genomic study in the Valencia Region (Spain) from January 2016 to December 2016. Short-read sequencing data of MTBC isolates collected from January 2014 to December 2016 were previously generated and published on the European Nucleotide Archive (ENA) in the BioProjects PRJEB29604, PRJEB38719, PRJEB65844, PRJEB89397, and PRJEB70424. Isolates from January 2017 to December 2019 were newly sequenced following the methodology described in Cancino-Muñoz et al., 2022^16^, and can be found under BioProject accession PRJEB89456. Only one representative sample from each patient was included in all the analyses except for the within-host diversity analysis (see below). Serial isolates for this analysis were deposited on BioProjects accessions PRJEB89397 and PRJEB65844.

### High-quality, high-integrity genomic DNA extraction and long-read sequencing

To determine if available genomic DNA samples from 2016 were suitable for long-read sequencing, we analyzed the requirements established by the manufacturer. We measured the DNA concentration using the Qubit® 3.0 Fluorometer (Thermo Fisher Scientific Inc., Waltham, Massachusetts), the DNA Integrity Number (DIN) using the TapeStation system (Agilent Technologies Inc., Santa Clara, CA), the A260/A280 and A260/A230 ratios using Tecan’s NanoQuant Plate® (Tecan Trading AG, Switzerland), and the presence of RNA by comparing the concentration of DNA detected by Qubit and NanoQuant. Isolates for which the genomic DNA did not meet the requirements were grown in solid media from the frozen culture for three weeks at 37°C. We optimized the standard CTAB-based DNA extraction protocol to obtain high-quality, high-molecular-weight genomic DNA. Finally, all the DNA samples received RNAse treatment (1 hour at 37°C) to remove contaminant RNA and were purified using magnetic beads (0.45x ratio) to select the longest DNA fragments. Our step-by-step optimized protocol can be found in protocols.io. Long-read sequencing libraries were generated using the SMRTbell® Express Template Prep Kit 2.0 (Pacific Biosciences of California Inc., Menlo Park, CA), following the manufacturer’s protocol. PacBio HiFi sequencing was performed using the Sequel II (SQII) platform. HiFi reads were obtained using the Circular Consensus Sequencing (CCS) mode on the SMRT Link Software (Pacific Biosciences of California Inc., Menlo Park, CA).

### Assessment of long-read HiFi sequencing

Quality control of HiFi reads was performed using a customized bash script to determine basic statistics and LongQC (v1.2.0b)^17^. Before *de novo* assembly, long reads were classified by taxonomy with Kraken (v0.10.5)^18^, and those not belonging to the MTBC were removed with KrakenTools (v1.2)^19^ for decontamination. HiFi reads containing the remaining PacBio adapter sequences were removed with HifiAdapter (v2.0.0)^20^.

Additionally, we determined the single mismatch rate per base in HiFi reads. Long reads from each isolate were aligned to the inferred MTBC most likely common ancestor (MTBCA)^21^ reference genome with minimap2 (v2.26)^22^. Unmapped reads and secondary and supplementary alignments were discarded. The number of single mismatches per read was obtained using the pysam^23^ module through a customized Python script^24^. Finally, the error rate per base was calculated by normalizing the number of mismatches to the number of aligned bases per read and then averaging across all reads.

### *De novo* assembly and manual curation

Assemblies from HiFi reads were built using Flye (v2.9.2)^25^ with two polishing iterations and circularized using Circlator (v1.5.5)^26^. To identify misassemblies caused by collapsed or expanded repeats, long reads were mapped to the assemblies with pbmm2 (v1.13)^27^, and genome coverage was plotted using the package ggplot2^28^ from R (v4.2.2)^29^. Regions whose coverage was significantly higher or lower than average were visually inspected using Artemis (v18.2.0)^30^, and assembly errors were manually corrected. Finally, since single-base substitutions and short indel errors can occur during the generation of draft genome assemblies, final consensus sequences were polished using their long reads. The polishing process involved three steps: i) mapping long reads to the assembly with pbmm2 (v1.13)^27^; ii) calling variants with freebayes (v1.3.6)^31^ and normalizing using vt (v0.57721)^32^; and iii) correcting positions in the assembly with the consensus allele (frequency above 0.5) using a customized Python script^24^. Finally, genome annotation was lifted over from the H37Rv reference genome (GenBank accession: AL123456.3) to each assembly, employing liftoff (v1.6.3)^33^ with -copies and -overlap 0.2 parameters and blastn (v2.12.0+)^34^ for complicated genomic regions.

### Quality control of genome assemblies

We evaluated three dimensions for a comprehensive assembly quality assessment: contiguity, completeness, and correctness^35^. The methods used in this analysis are reference-free because distinguishing between assembly errors and genuine biological differences is challenging and can lead to misinterpretation of results^36^.

Contiguity reflects how well the assembled sequence covers the underlying genome and was assessed using the following metrics: number of contigs, total length, circularization, and N50.

Completeness evaluates the gene content of the assembly and was measured with: i) BUSCO (v5.5.0)^37^ to identify the presence or absence of highly conserved genes, ii) IDEEL^38^ to determine the proportion of predicted proteins that are at least 95% the length of their best-matching known protein in a database, and iii) Long-read mapping coverage and the proportion of mapped/unmapped reads, iv) Merqury (v1.3)^39^ to calculate k-mer completeness using short reads, v) Prodigal (v2.6.3)^40^, to measure the average length of predicted proteins, and vi) QUAST (v.5.2.0)^41^ to obtain the GC content.

Correctness concerns the accuracy of each base pair in the assembly and how accurately contigs represent the genome sequence. To assess this feature, we measured: i) the concordance between the assembly and short-read data by mapping short reads against the assembly to calculate horizontal coverage and proportion of mapped/unmapped reads and detect small mismatches with Freebayes (v1.3.6)^31^, ii) the presence of high frequency (above 0.9) misassemblies detected by Sniffles2 (v2.3.3)^42^ using long reads, and iii) the assembly quality value (QV) by Merqury (v1.3)^39^, which measures the base-level accuracy using short-read k-mers.

### Multiple Genome Alignment from Complete Genomes

We obtained a Multiple Genome Alignment (MGA) that included 216 complete genomes and the MTBCA reference genome using Minigraph-Cactus (v2.8.4)^43^ with default parameters. The final MGA, originally in HAL (Hierarchical Alignment) format, was converted to MAF (Multiple Alignment Format) format using cactus-hal2maf with the parameters dupeMode single and noAncestors from HAL tools (v2.3)^44^. Synteny blocks in the MGA were identified using maf2synteny (v1.2)^45^.

The complexity of MGA algorithms is known to affect alignment accuracy. To minimize the risk of false positive variant calls, we refined the initial MGA by masking ambiguous positions. These were defined as positions with single variants not supported by at least one of two pairwise genome alignment approaches (nucmer^46^ or minimap2^22^). First, we obtained pairwise variant calls between the MTBC ancestor reference genome and each sample from the MGA in MAF format using a customized Python script^24^. In parallel, pairwise alignment and variant calling were performed between the MTBC ancestor reference genome and each complete genome with two different tools: i) nucmer (v4.0.0rc1)^46^ with the maxmatch parameter and dnadiff (v1.3)^47^ for variant calling. ii) minimap2 (v2.26)^22^ with the --asm20 parameter and paftools (v2.26)^22^ for variant calling. A position was masked in a particular sample of the MGA when it harboured a variant only detected in the global alignment. If the number of samples with a variant in a specific position differed by more than 10% between the three approaches, the position was masked in all the samples of the MGA.

Additionally, positions with two or more alleles at frequencies between 0.1 and 0.89 were masked in the MGA, ensuring that only fixed variants were included in the downstream analyses.

### Reference-mapping genomic analysis using short reads and phylogenetic analysis

We conducted a standard bioinformatic analysis to analyze short-read data, which included read trimming and filtering, read decontamination, mapping to a reference genome, and variant calling. We applied a validated pipeline^48^ that performs comparably to those developed by major Public Health TB reference laboratories. The workflow overview is: i) Mapping MTBC short reads to the MTBCA reference genome using BWA mem (v0.7.10)^49^, ii) Variant calling with VarScan2 (v2.3.7)^50^ and GATK HaplotypeCaller (v3.8)^51^. Single nucleotide polymorphisms (SNPs) called with high confidence are categorized into: fixed SNPs with allele frequency (AF) equal to or above 0.9, and non-fixed SNPs with AF between 0.1 and 0.89, and iii) Annotation of variants using SnpEff (v4.1)^51,52^ with H37Rv as reference (GenBank accession: AL123456.3) and masking of repetitive regions such as PE/PPE family and mobile elements. Lineage typing was performed by comparing SNPs with a catalog of phylogenetic marker positions^48^. Drug resistance profiles were determined by identifying mutations associated with resistance, as listed in the World Health Organization (WHO) catalog^53^. Specific parameters for each step can be found at the following repository: http://tgu.ibv.csic.es/?page_id=1794.

The reconstruction of transmission networks using short reads was performed in two steps: i) Construction of multiple sequence alignment (MSA) files with concatenated fixed SNPs and discarding invariant positions with snp-sites (v2.5.1)^54^, and ii) Construction of the genomic network with the MSA files applying a median-joining network inference method implemented in PopArt Software (v1.7)^55^.

### Comparison between complete genomes and short-read data

Pairwise genetic distances between all pairs of samples were calculated from the refined MGA by applying a customized Python script^24^. As for short-read data, pairwise genetic distances were calculated with the R ape^56^ package from a MSA file with concatenated SNPs in unmasked genomic regions. In both cases, only fixed SNPs were considered. The correlation between pairwise genetic distances from both short-read and complete genome data was determined using the Spearman rank coefficient.

Additionally, we compared the phylogenetic topology between the short-read derived and the complete genome derived phylogeny. Phylogenies were inferred using the maximum likelihood method with the GTR evolutionary model in IQ-TREE2 (v2.2.5)^57^, applied to both the MSA from the short-read data and the refined MGA from the complete genome data. We then compared these two phylogenies using the Shimodaira-Hasegawa (SH) test in IQ-TREE.

After confirming that both trees were topologically congruent, we assessed the accumulation of mutations in each branch of the phylogeny (branch lengths), comparing the short-read MSA alignment to the short-read MGA alignment. We then used the tree derived from short reads as a reference to map the polymorphic positions from the complete genome MGA using TreeTime (v0.11.4)^58^. We correlated the branch lengths derived from the short-read alignment to the branch lengths from the complete genome alignment.

### Estimation of evolutionary rates

Bayesian phylogenetic analyses were conducted using BEAST (v2.7.7)^59^ to estimate evolutionary rates from *M. tuberculosis* lineage 4 samples (n = 210), using short-read sequences and complete genomes independently. The site model was set to GTR+Γ4 with empirical base frequencies in BEAUti. A strict molecular clock was applied with a LogNormal prior on the evolutionary rate (M = 8.0, S = 1.2, in real space), and an initial value of 4.6 × 10⁻⁸ substitutions/site/year, following Bos et al. (2014)^60^. A coalescent constant population model was selected for the tree prior, with a 1/x prior distribution on population size.

As the dataset was homochronous, an ancient DNA sequence from a 17th-century Bishop (1679 AD)^61^ was included to calibrate the molecular clock in the short-read sequences analysis. For the complete genome analysis, the posterior time to the most recent common ancestor (tMRCA) from the short reads analysis was used as a prior, modeled as a normal distribution under three scenarios to assess robustness: i) mean = 200, SD = 150; ii) mean = 300, SD = 100; and iii) mean = 316, SD = 20. Ascertainment bias was corrected by defining invariant positions in the XML file, as recommended^62^.

For each dataset, three independent Markov chain Monte Carlo (MCMC) runs were conducted with chains of length 50 million and 10 million for each short-read and complete genome scenario, respectively. Convergence and mixing were assessed using Tracer (v1.6)^63^, with effective sample size (ESS) values >200 considered acceptable following a 10–20% burn-in. Posterior distributions of the evolutionary rate from each run were visualized using the R packages ggplot2^28^, tidyverse^64^, and ggridges^65^.

### Genetic diversity across the complete MTBC genome

To describe genomic diversity at global and local scales, we measured two parameters: i) the number of segregating sites (S) to assess global genetic diversity accumulated since the emergence of the MTBC by comparing to the MTBC common recent ancestor genome, and ii) nucleotide diversity (π) to describe intrapopulation genetic diversity accumulated between extant strains of the MTBC. A sliding window analysis was performed across the refined MGA, with segments of 200 bp at 1 bp intervals, using Variscan (v2.0.5)^66^. Regions with sequence coverage below 85% were excluded, and only windows with a size greater than one standard deviation below the mean were included. The number of segregating sites was normalized by window size to consider large gaps in the MGA. Regions with significantly higher diversity were identified by calculating the z-score for normalized S and π in each window. Then, genes and intergenic regions in windows with a z-score greater than five were annotated. The distribution of both the number of segregating sites and nucleotide diversity was plotted using the ggplot2^28^ package included in R (v4.2.2)^29^.

In addition, we performed a gene-enrichment analysis to identify Clusters of Orthologous Groups (COGs) that were overrepresented in our list of genes with higher diversity. Each gene was assigned to a specific MTBC COG category^67^. We performed a Fisher’s exact test for each COG, and adjusted p-values were obtained by applying the Benjamini-Hochberg correction.

### In-depth analysis of the *pe/ppe* gene family

As *pe/ppe* genes are highly complex regions of the MTBC genome, we analyzed them separately. The single *M. bovis* strain was excluded from this analysis. We extracted sequences of *pe/ppe* genes from each complete genome based on the gene coordinates in the H37Rv annotation (Genbank: AL123456.3). We then constructed a MSA of each gene with MACSE (v2.07)^68,69^, which aligns protein-coding sequences accounting for frameshifts, using the parameters -gc_def 11, -fs_term 20.0, and -fs 40.0. Sequences with at least 80% coverage and identity were included to reliably identify homologous positions for generating alignments. MSAs were curated with Aliview and manually corrected when needed.

To describe the gene conversion events within the *pe/ppe* family, we used blastn (v2.12.0+)^34^ to align the targeted region against the complete genomes and the MTBCA reference genome. We identified a gene conversion event when obtaining two complete 100% identity hits against the complete genome, but only one against the MTBCA reference genome, which pinpointed the ‘donor’ sequence location. Additionally, we used Gubbins (v3.4)^70^ to illustrate the frequency of these events across the gene family. We concatenated all the MSAs into a single file using FastaCon^71^ and plotted the recombination events of the different genes using Gubbins^70^ with IQTREE, GTR model and 1000 bootstrap replicates for tree building and a maximum window size of 4000bp to avoid overlapping.

Nucleotide diversity was measured for each gene using Variscan (v2.0.5)^66^. The analysis included all genes, even those not identified in all samples. Only positions in at least 85% of sequences in the MSA were considered.

To study structural variation in the locus *ppe38-71*, we identified *ppe38*, *esxX*, *esxY*, and *ppe71* using blastn from their annotation in the CDC1551 reference genome (Genbank: NC_002755.2, mt2419-22) as suggested by Ates, 2020^72^, and the GenBank X17348.1 sequence for the mobile element IS6110.

### Evolutionary analysis of T cell epitopes of the PE/PPE antigens

We accessed the Immune Epitope Database (IEDB)^73^ (https://www.iedb.org/) to obtain all the T cell epitope-encoding sequences of the *pe/ppe* gene family on January 16, 2025. We applied the following search parameters: linear peptide, organism *Mycobacterium tuberculosis* (ID:1773), T cell assays, any MHC Restriction, human host, and infectious disease. We retrieved 264 epitopes of the *pe/ppe* family. We performed a tblastn (v2.12.0+)^34^ against the H37Rv reference genome to identify the epitope’s genomic coordinates. After removing those with no hits found (40), identity below 80% (4), hits in a different gene from the one indicated in the database (4), and duplicates (6), 210 epitope sequences remained. Of those, 124 overlapping sequences were merged. Finally, we obtained 115 epitopes belonging to 61 *pe*/*ppe* genes.

We adopted the methodology of Comas *et al*. 2010^5^ to assess the evolutionary signatures in the *pe/ppe* genes. First, we established five categories and generated a concatenated MSA for each one: epitopes and non-epitope sequences from antigens, complete antigens, and essential and non-essential genes. We then used SNAP^74^, based on the Nei and Gojobori method^75^, to calculate the number of synonymous and non-synonymous substitutions in the five concatenates, comparing each sequence with the MTB ancestor reference genome. Finally, we obtained the medians of the dN/dS ratio for each group and compared them using the Wilcoxon signed-rank test. Additionally, we conducted the same analysis for epitopes, non-epitopes, and complete antigens, after removing gene conversion events^76^.

### Pairwise analysis of genomically closed samples using complete genomes

We used short-read data to identify MTBC pairs at genetic distances of up to 20 fixed SNPs, thus capturing recent and older transmission events according to conventional short-read pipelines. For each pair of closely related samples, pairwise alignments and variant calling using complete genomes were performed with the varifier (v0.4.0)^77^ make_truth_vcf function, which uses dnadiff (v1.3)^47^ and minimap2 (v2.26)^22^ to build a consensus call. To detect potential false negative SNP calls, alignments were manually inspected. Positions with a non-fixed SNP were masked in each assembly. Only fixed indels (with AF above 0.9) in each assembly were included. Indels detected in homopolymer sequences (three or more nucleotides) were excluded, as long-read sequencing technologies are more error-prone in these regions, making it challenging to discern between sequencing errors and real variation.

### Multiple genome alignments to reconstruct transmission networks

Individual MGAs were constructed using progressiveMauve (v2.4.0)^78^ with default parameters, for each transmission cluster previously identified by short-read data, applying a 10-SNP threshold. Mutations were extracted from MGAs using the msa2vcf tool from jvarkit^79^ and validated with pairwise alignment data. Only the variants that met the criteria specified in the previous section were included in the network analysis. SNP and indel data were used to reconstruct transmission networks by applying the median-joining network inference method with PopArt Software (v1.7)^55^. Additionally, in one cluster, we identified a gene conversion event between two genes belonging to the *pe/ppe* family. This recombination event was considered one mutation event in the network’s construction.

Once we identified the indels between samples of the same cluster, we aimed to clarify whether they were medium-small indels or structural variations. We performed pairwise genome alignments between all pairs of samples in transmission using Nucdiff (v2.0.3)^80^ with default parameters. Additionally, to identify whether some duplications were insertions of the mobile element IS6110 (Genebank accession: X17348.1), we compared them using blastn.

### Assessment of variant calling from short-read sequencing data in masked regions

To evaluate the performance of variant calling from short-read mapping in regions systematically excluded from standard analyses, we assessed each masked position with an alternative genotype, using complete genomes as the ground truth. We detected ground truth SNPs by performing variant calling on the refined MGA, comparing the MTBCA reference genome with each complete genome using a customized Python script to parse the MAF file and identify variants relative to the reference^24^. Positions that were not validated during the refinement of the MGA for each sample were excluded from these measurements.

Variant calls from short-read data were obtained using the pipeline described earlier. In both complete genome and short-read mapping analyses, we focused exclusively on fixed SNPs located in regions discarded by the annotation filter. Agreement between the two methods was assessed by calculating recall (the proportion of ground truth variants correctly identified by short-read mapping), precision (the proportion of true variants among the variants detected by short-read mapping), and F1 score (harmonic mean of recall and precision) for each polymorphic position detected in at least 10 samples.

### Cluster-specific reference genomes for enhancing epidemiology analyses

The standard bioinformatic and downstream phylogenetic analyses were performed using the pipeline described above on short-read data from all MTBC isolates collected in the Valencia Region from 2014 to 2019. We selected the transmission clusters (applying a 10-SNP threshold) with at least one isolate from our dataset of complete genomes, allowing us to designate one of them as a cluster-specific reference genome. If more than one complete genome was available, we chose the one from the patient who was diagnosed first. We then reanalyzed each transmission group twice, using a different reference genome for each analysis: one with the cluster-specific complete genome and the other with the MTBCA as reference genomes. We then compared basic indicators of short-read mapping quality. Additionally, both analyses were performed using two filters: i) a standard, annotation-based filter from Comas et al. 2010^5^, and ii) a refined version adapted from Marin et al. 2022^4^ that keeps regions with good mappability and base recall. Both filters are available in our repository: https://gitlab.com/tbgenomicsunit/Publications_resources.

Non-fixed SNPs, defined as calls with AF between 0.1 and 0.89 (inclusive), were analyzed in detail by comparing their distribution between the different reference and filter approaches. The cluster-specific reference genome approach allowed us to incorporate non-fixed variants that were fixed in another sample within the cluster into transmission analyses. To ensure the reliability of these non-fixed calls, we applied stricter criteria, excluding all supplementary alignments and reads without a proper pair mapped^81,82^ from the short-read alignment before the variant calling. As an indicator of improved epidemiological resolution, we assessed: i) clusters with changes in topology, and ii) clusters with changes in relative distances between samples but maintaining the topology.

### Patient-specific reference genomes to study within-host evolution

To study the impact of using a patient-specific reference genome for detecting within-host diversity, we analyzed five clinical cases from the Valencia Region in 2016. For each patient, we had a complete genome of the first isolate along with matched short-read data, as well as at least one additional isolate with short-read data obtained at least one week later. In one case, a third isolate was also available. We performed the previously described bioinformatic analysis using the MTBCA and a patient-specific complete genome as references. In addition, we conducted the analysis twice for each reference approach, using both the standard and the refined masking filters. We then identified non-fixed SNPs across all isolates and assessed the proportion of non-fixed calls detected by the common reference approach that was validated by the patient-specific reference approach.

## RESULTS

### A unique dataset of paired high-quality MTBC complete genomes and short-read data

Of 266 MTBC culture-positive isolates available in the Valencia Region of Spain from 2016, 216 (80%) were sequenced using the PacBio HiFi method. An average sequencing depth of 100.5x was achieved with a mean read length of 4,505 (see Supplementary Notes, Fig. S1, and Table S3 for more statistics). We achieved 212 genome assemblies closed in a single circular contig, and all genome assemblies were highly accurate and complete. A detailed description of parameters used to assess contiguity, completeness, and correctness can be found in Supplementary Notes, Extended Data Table 1 and Table S4. As the 216 assemblies demonstrated consistently high-quality metrics, they were all included in downstream analyses.

Short-read data were available for all 216 MTBC isolates, with a mean coverage depth of 130x. The lineage (L) distribution of the MTBC samples was as follows: 4 (2.0%) from L2, 2 (0.9%) from L3, and 209 (96.7%) from L4. In addition, one sample (0.4%) was identified as *M. bovis*. 194 (89.8%) isolates were susceptible to all first-line antituberculous drugs, and 22 (10.2%) isolates were resistant to at least one antibiotic (drug resistance profiles are detailed in Fig. S1 and Table S3).

### Complete genomes reveal greater genetic diversity and a higher evolutionary rate

To assess the performance of the complete genome approach in detecting SNPs, we measured the pairwise genetic distances between each pair of samples comparing complete genomes and short-read data. The two measures were strongly correlated (Spearman’s rho = 0.87, p < 2.2e-16), with complete genomes consistently yielding equal or greater pairwise genetic distances than those obtained by short-read mapping, showing an average of 303 and a median of 312 (-1 to 792) additional SNPs per pair (Fig. 1a). Of the additional SNPs identified, 88% (268) were in regions discarded by the short-read mapping approach, while 12% (35) were found in preserved regions. Moreover, the phylogeny of complete genomes was not significantly different than the short-read derived (SH test p ≈ 1), with a strong correlation between all branch lengths (Spearman’s ρ = 0.98, p < 2.2e-16). These results suggest that the additional variation was evenly distributed throughout the phylogeny, with no evidence of accumulation in specific branches.

Additionally, to quantify the additional variation, we estimated evolutionary rates for *M. tuberculosis* lineage 4 strains using both short reads and complete genomes (Fig. 1b,c). For the short-read sequences, the estimated median evolutionary rate was 5.14 × 10⁻⁸ substitutions/site/year, with a 95% highest posterior density (HPD) interval of 3.94 × 10⁻⁸ to 6.34 × 10⁻⁸. In contrast, the complete genome analyses yielded consistently higher rates across three prior scenarios: i) 7.84 × 10⁻⁸ [95% HPD: 6.47 × 10⁻⁸ – 9.50 × 10⁻⁸]; ii) 7.88 × 10⁻⁸ [6.96 × 10⁻⁸ – 8.93 × 10⁻⁸]; and iii) 7.60 × 10⁻⁸ [7.31 × 10⁻⁸ – 7.88 × 10⁻⁸]. Overall, evolutionary rate estimates derived from complete genome data were 1.5-fold higher than those from short reads.

**Figure 1.**
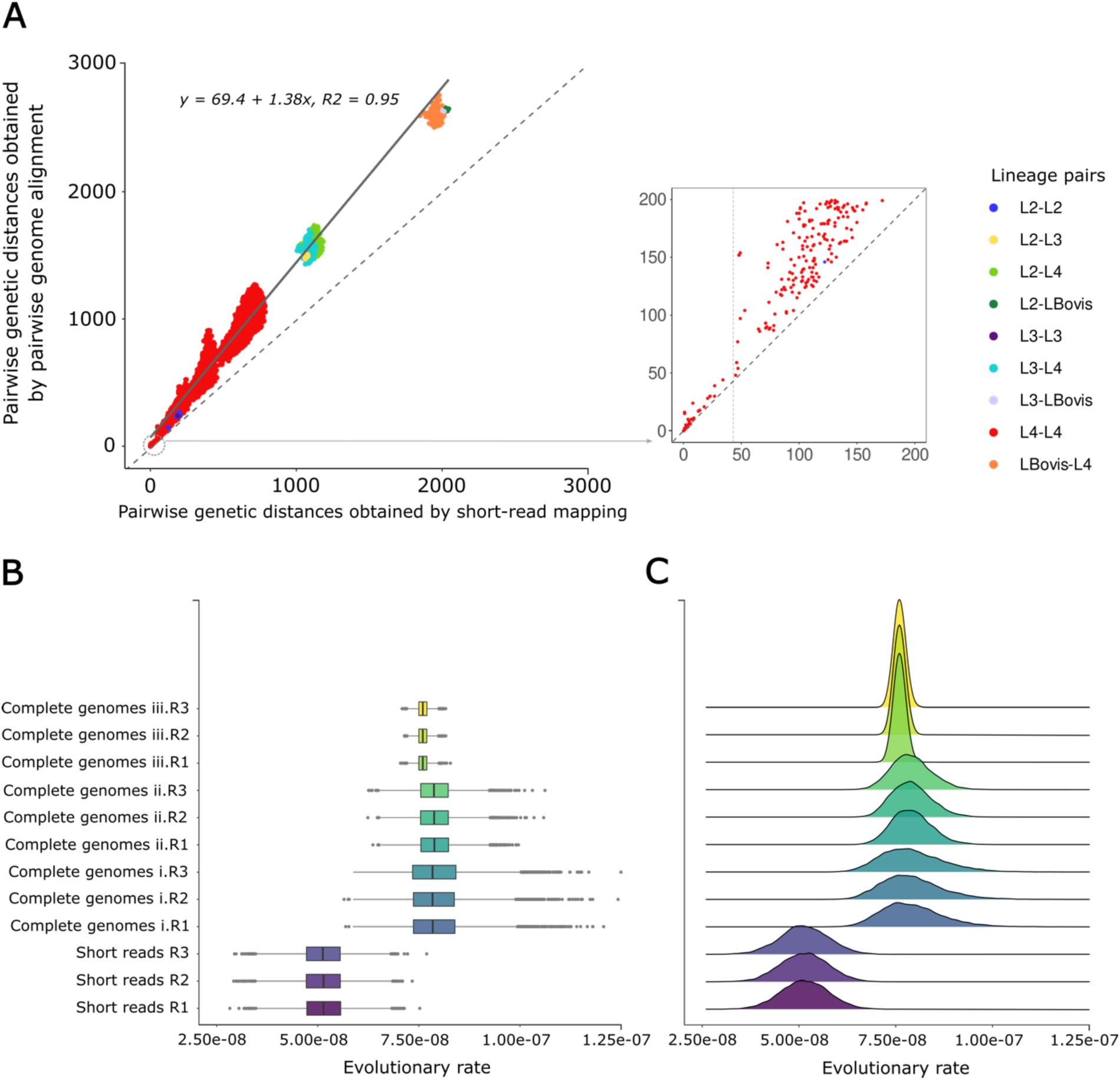
Comparison of pairwise genetic distances and evolutionary rates using short-read and complete genome data. (A) Comparison of pairwise genetic distances between MTBC isolates using short-read and complete genome data. Each point represents a pairwise comparison between the 216 MTBC isolates, with SNP distances obtained from short-read data on the x-axis and complete genomes on the y-axis. The dashed gray line represents the identity line (x = y), while the solid line indicates the linear regression. Notably, almost all comparisons fall above the identity line, indicating higher genetic distances obtained from complete genomes. Colors indicate the lineages of the strains involved in each pairwise comparison. The three differentiated dot clusters correspond to varying levels of divergence: the first includes comparisons between isolates of the same lineage, the second between human-adapted lineages, and the last between *M. bovis* and human-adapted lineages. This illustrates how the number of SNPs gained increased with the level of divergence between the isolates, thus, when comparing isolates from distinct lineages. The inset zooms in on the region with ≤200 SNPs in both approaches and the vertical dashed line marks the genetic distance obtained by short reads at which comparisons begin to scatter. It shows that pairs of isolates with closer genetic distance (based on short-read data) showed a substantially lower SNP count discrepancy that increased sharply when comparing more genetically distant isolates. (B, C) Comparison of evolutionary rates estimated with short-read and complete genome data for MTB lineage 4 isolates from this dataset. For short reads, only one prior was analyzed, while three different priors were analyzed for complete genome data to test the robustness of the estimates; three replicates were performed for each scenario (y-axis). The number of substitutions/site/year is indicated on the x-axis. For each replicate, two distributions are shown: a boxplot (B) and a density plot (C).

### Global-scale analysis of complete genomes unveils genetic diversity hotspots across the MTBC genome

From the pairwise genetic distance analysis, we found that 20% (4804/25865) of polymorphic sites were in regions not assessed by standard short-read approaches. To characterize the total MTBC diversity in our dataset, including those regions, we performed a sliding window analysis across the refined MGA. Segregating sites between complete genomes and the MTBCA reference genome were analyzed to assess genomic divergence since the emergence of a common ancestor. This analysis identified 94 high-diversity genomic regions, including 64 genes and 30 intergenic regions (Fig. 2a, Table S5). Based on the masking filter applied in the short-read mapping approach, 60% (55/94) were in masked segments, while 40% (38/94) were in preserved segments. A functional analysis identified the PE and PPE families as major contributors to diversity within our dataset (Fig. 2c and Table S1).

**Figure 2.**
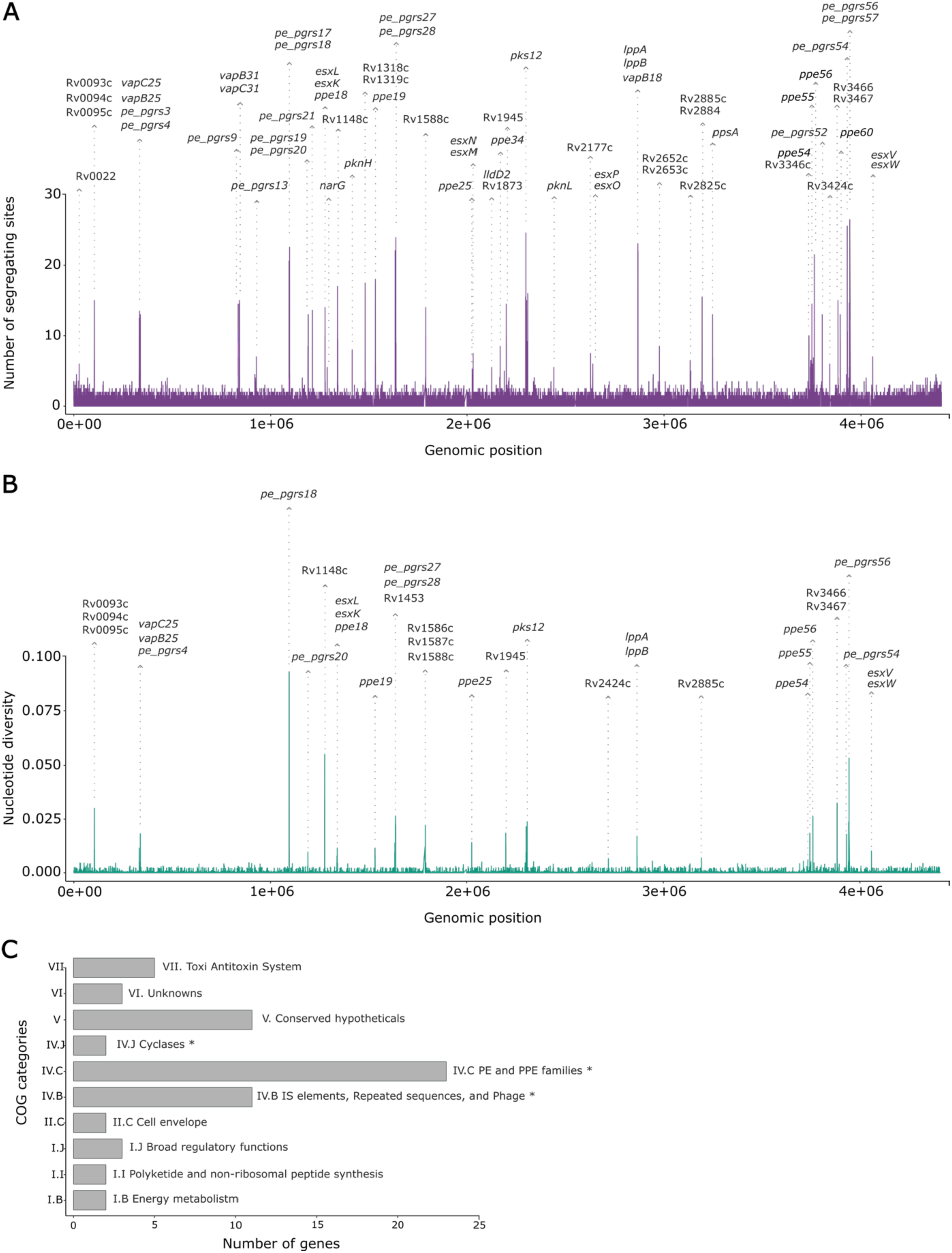
Genomic diversity of *Mycobacterium tuberculosis* complex. Sliding window analysis showing (A) the number of segregating sites and (B) the nucleotide diversity in blue across the refined MGA with 216 MTBC complete genomes. The window length was 200 bp with a step size of 1 bp. The x-axis indicates the position of the windows’ midpoint. (A) The analysis of segregating sites revealed that 60% (55/94) of diversity hotspots were in regions poorly characterized by short-read mapping, while 40% (38/94) were in preserved ones. (B) Nucleotide diversity was measured between strains in our population dataset, considering that most is contributed by Lineage 4. This analysis identified 52 high-diversity hotspots, 77% (40/52) were in discarded and 23% (12) in preserved regions (Fig. 2, Table S5). Genes with diversity hotspots were distributed across 10 MTBC COG (Clusters of Orthologous Groups) categories, displayed on the y-axis. The x-axis shows the number of genes per category. The functional enrichment analysis reveals significant overrepresentation in the following groups: IV.B IS elements, repeated sequences, and phages (adjusted p-value 6.86E-05), IV.J Cyclases (adjusted p-value 0.02), and IV.C PE and PPE families (adjusted p-value 2.58E-14); *, adjusted p-value < 0.01 (see also Table S1).

### Gene conversion is a main source of nucleotide diversity in the *pe/ppe* gene family

We generated MSAs for the 169 *pe/ppe* genes and measured their total nucleotide diversity. All genes were included in the analysis, with 65% of genes (109) present in all genomes (Table S6). As shown in Fig. 3a, nucleotide diversity varied widely across the family, ranging from highly conserved to hypervariable genes. Furthermore, genetic variation was unevenly distributed within individual genes (Fig. S2). We identified gene conversion as the primary driver of nucleotide diversity (Fig. 3b) and we accurately delineated these events, which were detected in 28 *pe/ppe* genes, displaying diverse SNP patterns in terms of frequency and extent (Fig. 3c). A detailed overview of these events is provided in Fig. 4.

In addition to gene conversion, we observed different structural variants in several *pe/ppe* genes. Polymorphisms in the locus *ppe38-71* are of particular interest, as it has been linked to the secretion of virulence factors such as PE_PGRS and PPE-MPTR proteins^72^. Inactivation of this locus by deletion or truncation of *ppe38* exhibits higher virulence patterns in mouse models of infection^72^. We found substantial structural diversity in the *ppe38-71* locus even within our dataset enriched in lineage 4: *ppe38* was intact in most genomes (169/216), altered in 44/216, and completely absent in 3/216 (Fig. 5). Unexpectedly, three samples had duplications of the locus. Most of these locus conformations are newly described, and their impact on virulence remains unknown.

**Figure 3.**
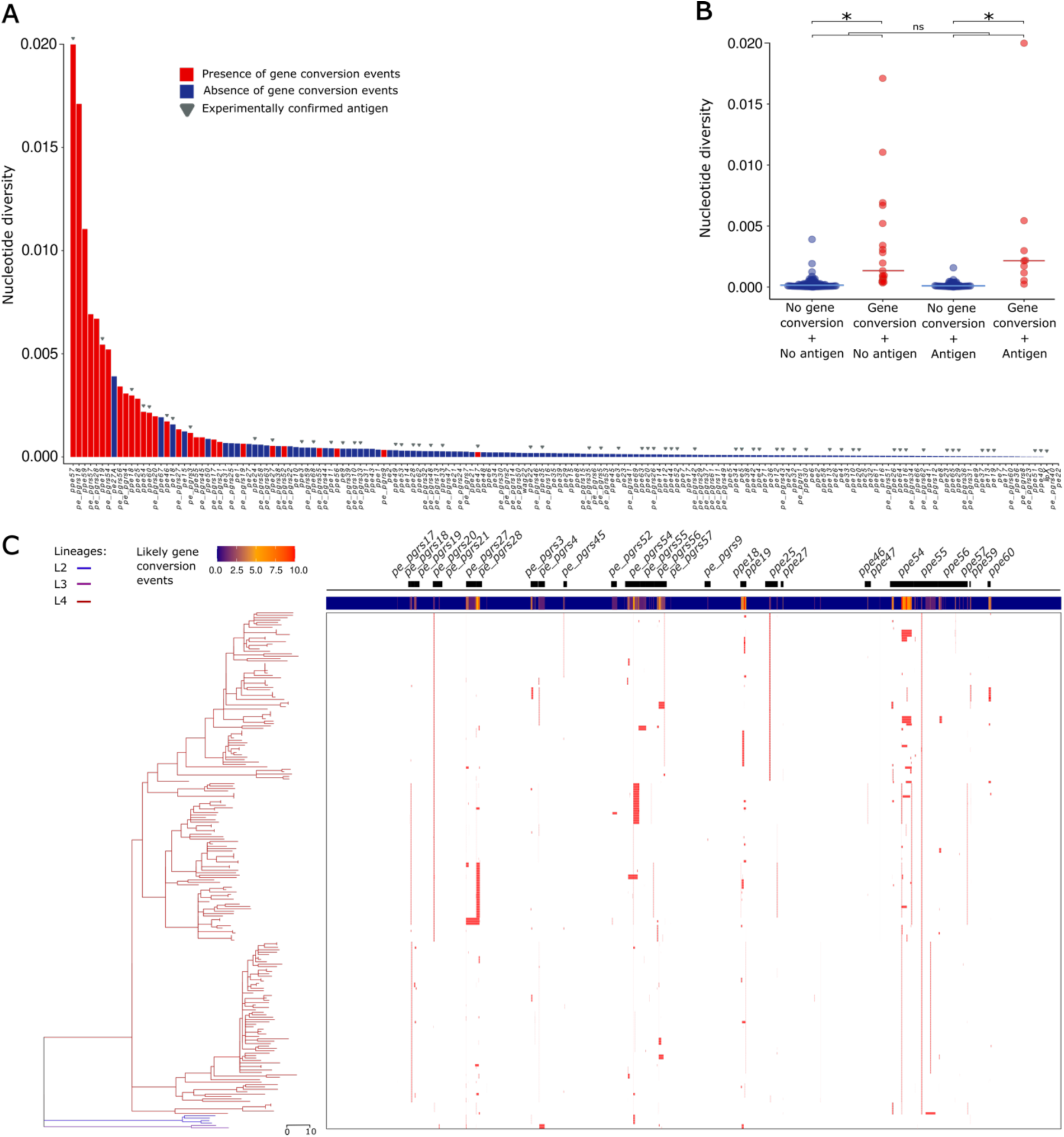
Genetic diversity of the *pe/ppe* gene family. (A) Distribution of nucleotide diversity of 169 *pe/ppe* genes. Genes with evidence of gene conversion events are in red, while the ones without evidence of gene conversion are in blue. Genes that encode a PE/PPE antigen are marked with a grey triangle. See Table S6 for specific data. (B) Distributions of nucleotide diversity of four *pe/ppe* gene categories. *Pe/ppe* genes were classified based on the presence of gene conversion events and whether they encode antigens. The distribution of nucleotide diversity was plotted for each category. Brackets indicate Wilcoxon Rank Sum tests of pairwise category comparisons; ns, not significant; *, p < 0.05. (C) Distribution of candidate gene conversion events in *pe/ppe* genes. The panel on the left shows the maximum-likelihood phylogeny built from the clonal frame of a concatenation of *pe/ppe* genes. The tree is colored according to the lineage of isolates, each corresponding to a row in the panel on the right. The annotation of each *pe/ppe* gene with evidence of gene conversion is shown at the top of the figure. The main panel displays the distribution of inferred recombination events in the *pe/ppe* genes of our dataset, which are colored red. A portion of those is later validated in our analysis. Fig. 4 displays the schematic representations of gene conversion events between and within the *pe/ppe* genes.

**Figure 4.**
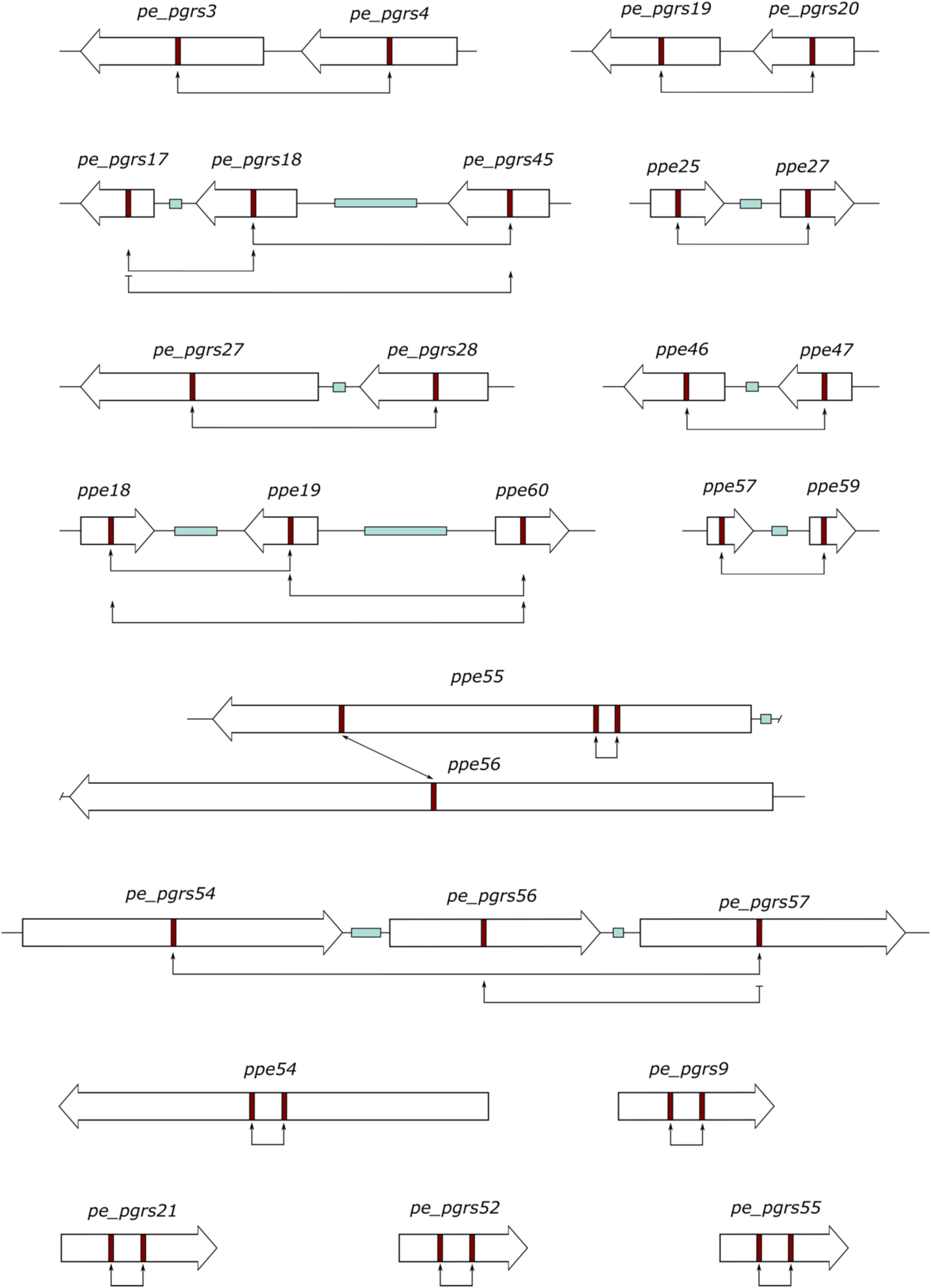
Schematic representations of gene conversion events between and within 28 *pe/ppe* genes. Each schematic represents a gene conversion event, illustrating the genes involved. Genes are depicted as thick arrows scaled to gene length and oriented according to their strand direction. Genes separated only by intergenic regions are connected by a thin line, while separation by additional loci is indicated by a blue bar. Intragenic events (within the same gene) are shown as two closely spaced red lines within the same arrow. Thin connecting arrows indicate the direction of gene conversion from donor to recipient genes. When arrows point in both directions, it indicates that those genes can function as both donors and recipients.

**Figure 5.**
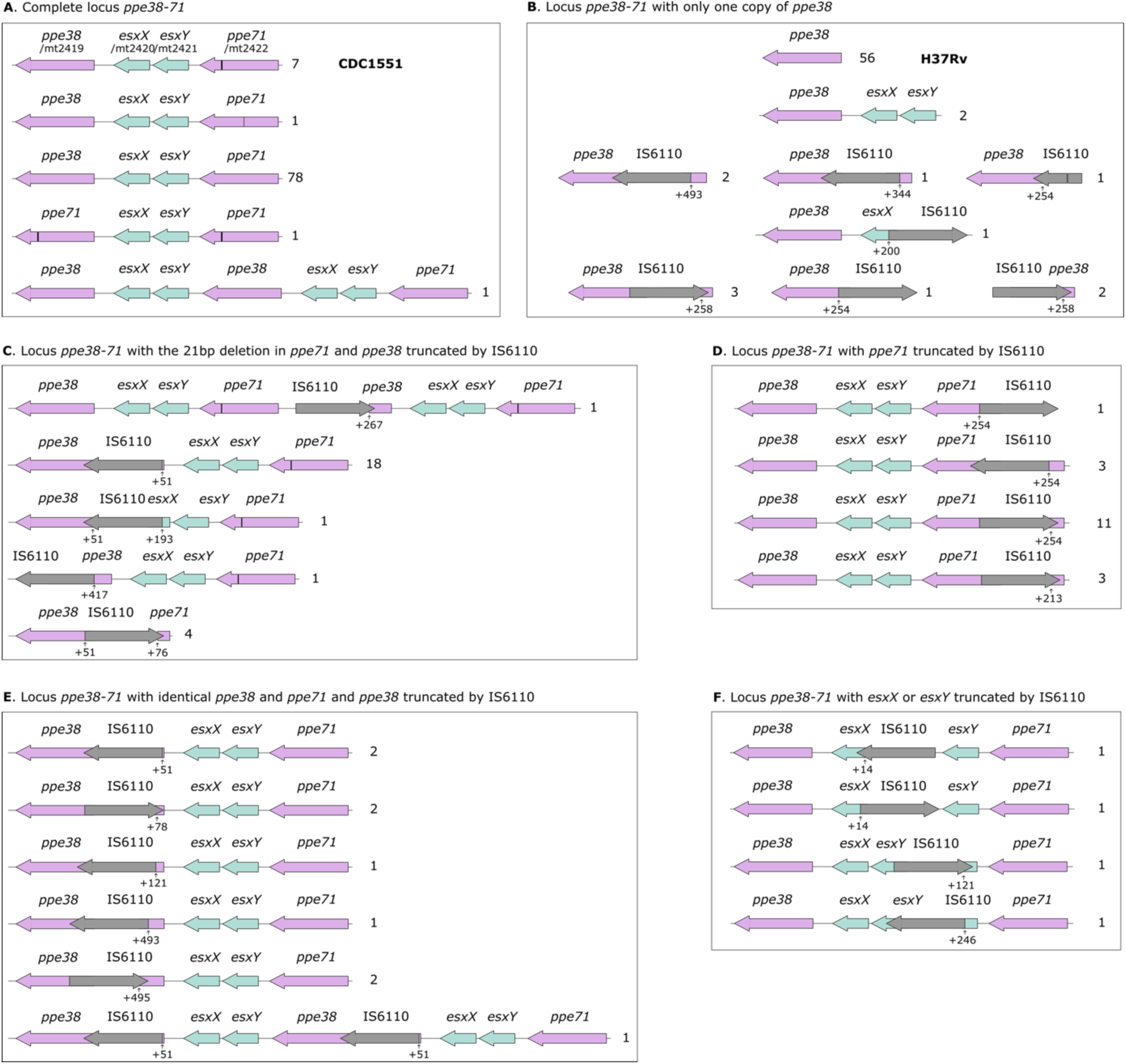
Schematic representation of polymorphisms of the *ppe38-71* locus. The *ppe38-71* locus includes *ppe38* (purple), *esxX* and *esxY* (blue), and *ppe71*, which may be identical to *ppe38* or contain a 21 bp deletion (indicated by a vertical black line). Due to a misannotation, the H37Rv reference genome (GenBank: AL123456.3) has only one copy of *ppe38*, but several H37Rv strains harbor the full functional locus. For this reason, we followed the annotation of the MTB CDC1550 reference genome (GenBank: NC_002755.2) to annotate this region in complete genomes, as recommended by Attes et al., 2020^72^. Multiple polymorphisms of locus *ppe38-71* were identified in the dataset and grouped into six main categories: a) complete locus; b) single copy of *ppe38* with various truncations due to IS6110; c) *ppe71* with a 21 bp deletion and *ppe38* truncated by IS6110; d) *ppe71* truncated by IS6110; e) *ppe71* identical to *ppe38*, with *ppe38* truncated by IS6110; f) complete locus with *esxX* or *esxY* truncated by IS6110. The number of samples corresponding to each genotype is indicated on the right. Three samples lacked the entire locus. Most of these loci conformations have unknown consequences for virulence since they are described for the first time. This is a salient example of how complete genomes can help to understand the evolution of traits such as virulence in MTB.

### T-cell epitopes of PE/PPE antigens are hyperconserved but can undergo gene conversion events

We investigated how intragenic variation affects *pe/ppe* antigen-encoding genes, focusing on T-cell epitopes, the part of antigens recognized by the immune system. We analyzed 61 experimentally confirmed human T-cell antigens encoded by *pe/ppe* genes (Table S7). PE/PPE antigens exhibited similar nucleotide diversity (p-value = 0.07, Wilcoxon signed-rank test) and were proportionally influenced by gene conversion events compared to other family members (Fig. 3b) (p-value > 0.05, Fisher’s exact test). Epitopes were significantly more conserved than non-epitope regions and full antigens (p-value < 0.01, Wilcoxon signed-rank test), with a median dN/dS of 0.17 versus 0.48 and 0.45, respectively (Fig. 6a). However, certain epitopes from *ppe57, ppe19*, and *ppe18* underwent gene conversion events, accumulating many non-synonymous substitutions (Fig. 6c). After excluding gene conversion events, we recalculated the dN/dS ratio and observed the same pattern of epitope hyperconservation (Fig. 6b), despite comparable levels of variation in epitope and non-epitope regions (6.8% vs. 6.1% segregating sites, p > 0.05, Fisher’s exact test). These are strong indicators of purifying selection acting in epitope sequences. Regarding structural variation in PE/PPE antigens, we identified small sequence duplications in PPE24 and PPE34 and large indels in PPE54 that did not affect epitopes. Structural variation primarily affecting non-epitope regions further supports the idea that epitopes tend to be under purifying selection.

**Figure 6.**
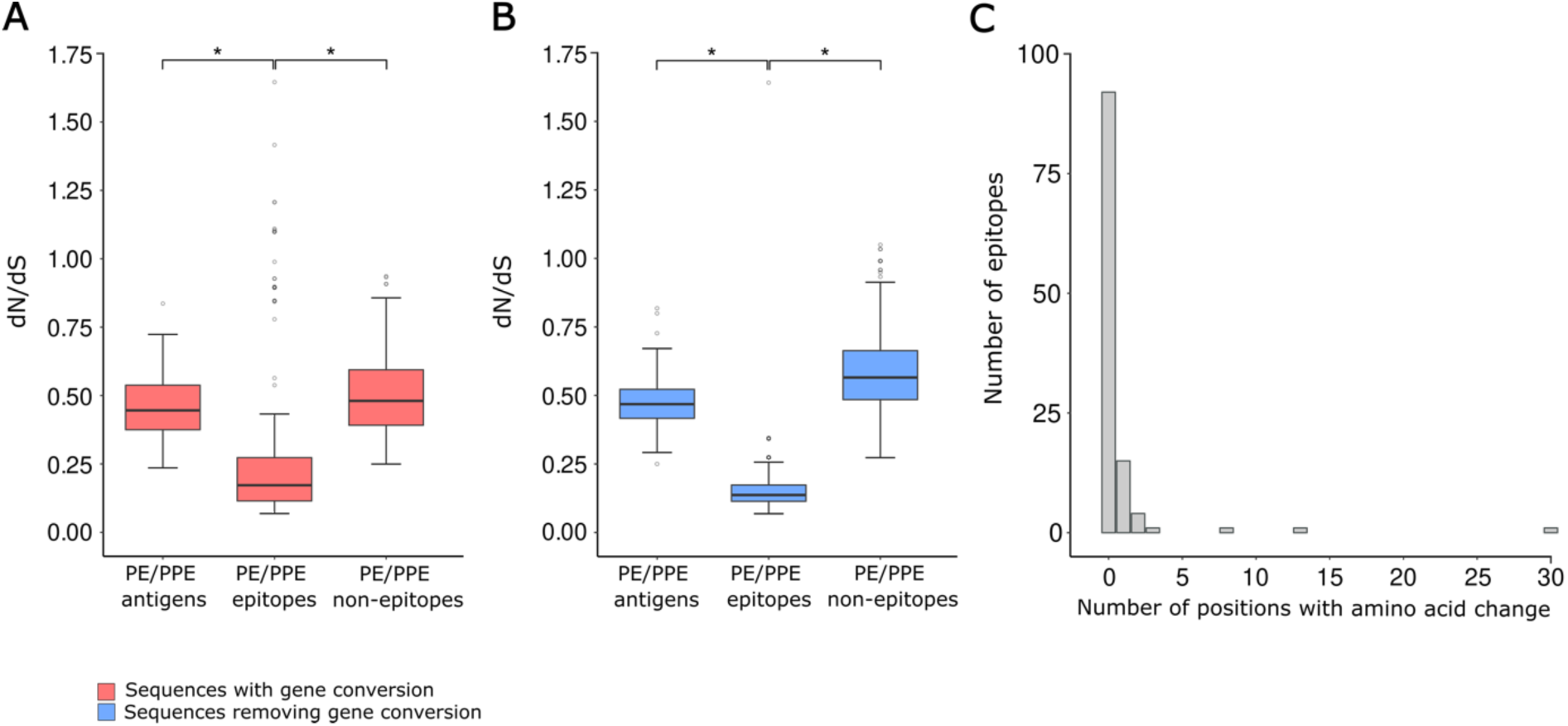
Evolutionary patterns of experimentally validated human T cell epitopes from PE/PPE antigens in *Mycobacterium tuberculosis*. (A) Comparison of dN/dS ratios across three different categories of PE/PPE antigens: 115 PE/PPE epitopes, the non-epitope regions of the same PE/PPE antigens, and 61 complete PE/PPE antigens. (B) Same comparison as the one shown in panel A, but analyzing the clonal frame of sequences exclusively (i.e., removing gene conversion events); *, p < 0.05. (C) Frequency distribution showing the PE/PPE epitopes grouped by the number of observed amino acid changes. Table S7 contains epitope sequences and additional genomic data.

### Complete genomes enhance genetic variant detection at a local scale and delineate recent transmission

We next focused on recently diverged isolates, analyzing pairs of isolates with a genetic distance of up to 20 SNPs, as determined by short-read data. Fifty-two pairs from 65 closely related isolates were examined in detail (Fig. 7a, Table S8). On average, 2.5 SNPs per pair were gained, with a median of 1 (-1 to 16) SNP and 2 mutations (SNPs and indels). Upon manual inspection, we identified one SNP, detected by short-read mapping but not complete genomes, as a false positive. In 60% (31) pairs, additional fixed SNPs were identified from the complete genome data, and fixed indels were found in 46% (24) pairs. Overall, additional variation was detected in 70% (36) pairs.

Eighty-eight percent (50/57) of the non-redundant SNPs detected only by complete genomes were in masked regions (Fig. 7b), with the *pe/ppe* gene family overrepresented as in the global analysis. The remaining 12% (7/57) SNPs were found in unmasked regions but were missed by the short-read mapping analysis (Supplementary Notes). Next, we investigated whether masked variants could have been retained in the short-read mapping approach. A detailed list with 1,081 polymorphic positions is provided in Table S9. F1 score distributions highlighted the value of masking individual positions rather than entire regions (also see Supplementary Notes).

**Figure 7.**
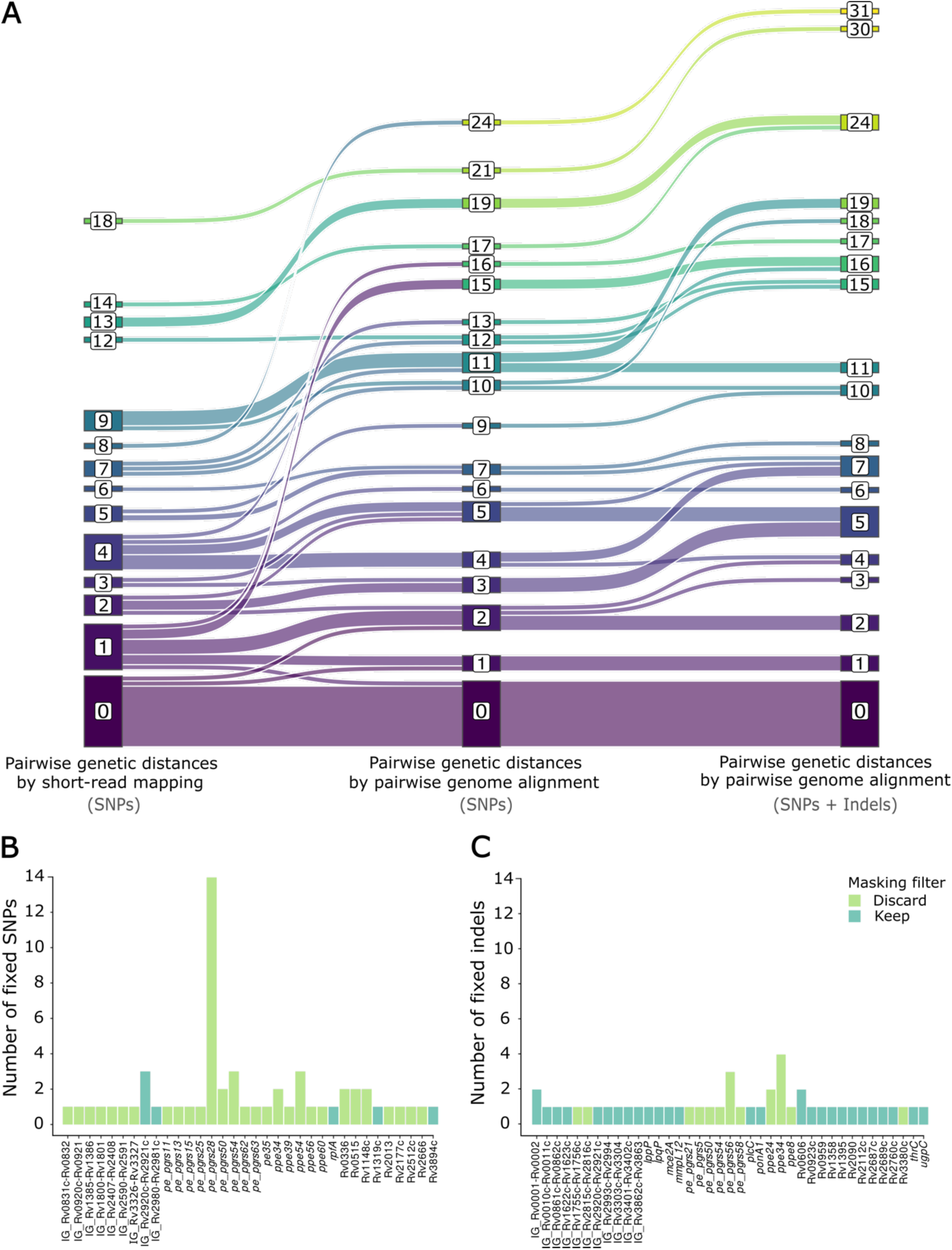
Gain of fixed mutations between closely related samples using complete genomes and their location in the genome. Pairwise genetic distances between samples at ≤20 SNPs according to short-read data were calculated by three approaches that included: i) short-read SNP data, ii) complete genome SNP data, and iii) complete genome SNP and indel data. The Sankey diagram (A) shows the increase in pairwise genetic distances when moving from short-read mapping to complete genome comparisons. Notably, 12 out of 14 pairs with 0 SNP distance remained unchanged, indicating that 0-SNP distances from short-read sequences are generally also reliable markers of very recent transmission. On the contrary, at 5 and 10 SNP thresholds, most pairs gain diversity considering only SNPs: 47% (18/38) and 57% (27/47) respectively; and when incorporating indels: 58% (22/38) and 66% (31/47) respectively. Furthermore, four pairs showed a substantially larger increase in SNPs (15 on average). A deeper investigation revealed that one isolate, common to all four pairs, underwent a gene conversion event between the *pe_pgrs27* and *pe_pgrs28* genes. (B) and (C) display the frequency and genomic distribution of additional SNPs and indels, respectively, detected only by complete genome data.

### Higher genomic transmission network resolution enabled by complete genomes over short-read mapping

We identified 24 transmission clusters based on a 10-SNP threshold using the short-read mapping approach, with cluster sizes ranging from 2 to 5 cases (Table S10). We then reconstructed the same clusters using complete genomes, analyzing SNPs, and SNPs combined with indels separately. Transmission network diagrams for all clusters and approaches are provided in Fig. S3. Complete genome analysis revealed more information in 62% (15/24) of the clusters, while 38% (9/24) remained unchanged. Among the clusters with differences, 21% (5/24) displayed a different topology, 8% (2/24) showed changes in edge lengths, altering the relative distances but maintaining the underlying topology, and 33% (8) only exhibited changes in edge lengths while maintaining the same relative distances and underlying topology. Regarding distances between samples in the network, 68% (32) of links showed changes, with an average of 2.5 mutations gained per link and a median of 2 mutations (-1 to 11), corroborating the pairwise distance analysis above.

We detected indels in 25% (6/24) of transmission clusters, with varying lengths and structural characteristics. The most common were medium-small indels, present in 8 isolates. Other structural variants included: 3 tandem duplications, 3 collapsed tandem repeats, 2 duplications, and 6 insertions of the IS6110. Additionally, we identified a gene conversion event from *pe_pgrs27* to *pe_pgrs28* that affected isolate G1823 in Cluster 10 (Fig. S3). It involved 14 additional SNPs that were included as a single mutation event in the analysis.

### Cluster-specific reference genomes allow for the incorporation of non-fixed variants into transmission analyses

To evaluate the impact of using a closer reference genome instead of a common reference, we analyzed short-read sequencing data from 372 MTB isolates belonging to 77 clusters (ranging from 2 to 21 samples) collected in the Valencia Region between 2014 and 2019 (Table S11). The comparison of both approaches demonstrated that using a closer reference genome provided greater reliability, particularly during the mapping step (see Supplementary Notes for details).

Notably, and linked to the next section, mapping to a cluster-specific reference genome revealed a sharp reduction of non-fixed variation compared to our conventional pipeline relying on a common reference genome. We observed a ten-fold reduction in non-fixed SNPs (Wilcoxon Signed-Rank Test, *p* < 2.2 × 10⁻¹⁶) (Fig. 8a), which resulted in a five-fold reduction when compared to using the refined masking filter with the common reference approach (Wilcoxon Signed-Rank Test, *p* < 2.2 × 10⁻¹⁶). The ability to accurately detect genuine non-fixed variant calls using a closer reference genome enabled the incorporation of this source of variation into the reconstruction of 40% (31/77) transmission clusters. We were able to enhance the epidemiological resolution of 23% (18/77) clusters (example shown in Fig. 8e).

### Patient-specific reference genomes reveal genuine within-host variation in serial isolates

We next explored whether mapping short-read data to a patient-specific reference genome also enhances non-fixed variant calling in patients with serial isolates. We analyzed five cases for which a complete genome and matched short-read data were available, along with at least one additional isolate sequenced by short-read technology (Table S11). Our results showed that 82% (524/640) of non-fixed variants detected when mapping short reads to a common reference and applying the standard masking filter were false positives. A median of 5.1% (0-63%) non-fixed SNPs per patient were validated by the patient-specific reference approach (Fig. 8b, Table S2). A refined filter partially ameliorates the issue of false positives. Still, 69.7% (276/396) of non-fixed SNPs were false positives, and a median of 10.8% (0-78%) of non-fixed SNPs was validated (Wilcoxon Signed-Rank Test, *p* = 0.04) (Fig. 8b). This filtering effect was not observed when using the patient-specific reference genome (Wilcoxon Signed-Rank Test, *p =* 0.7); in fact, some non-fixed SNPs located in previously masked regions were recovered, suggesting that masking filters may not be necessary for this approach. These findings have major implications for interpreting previously published studies analyzing within-host variation.

**Figure 8.**
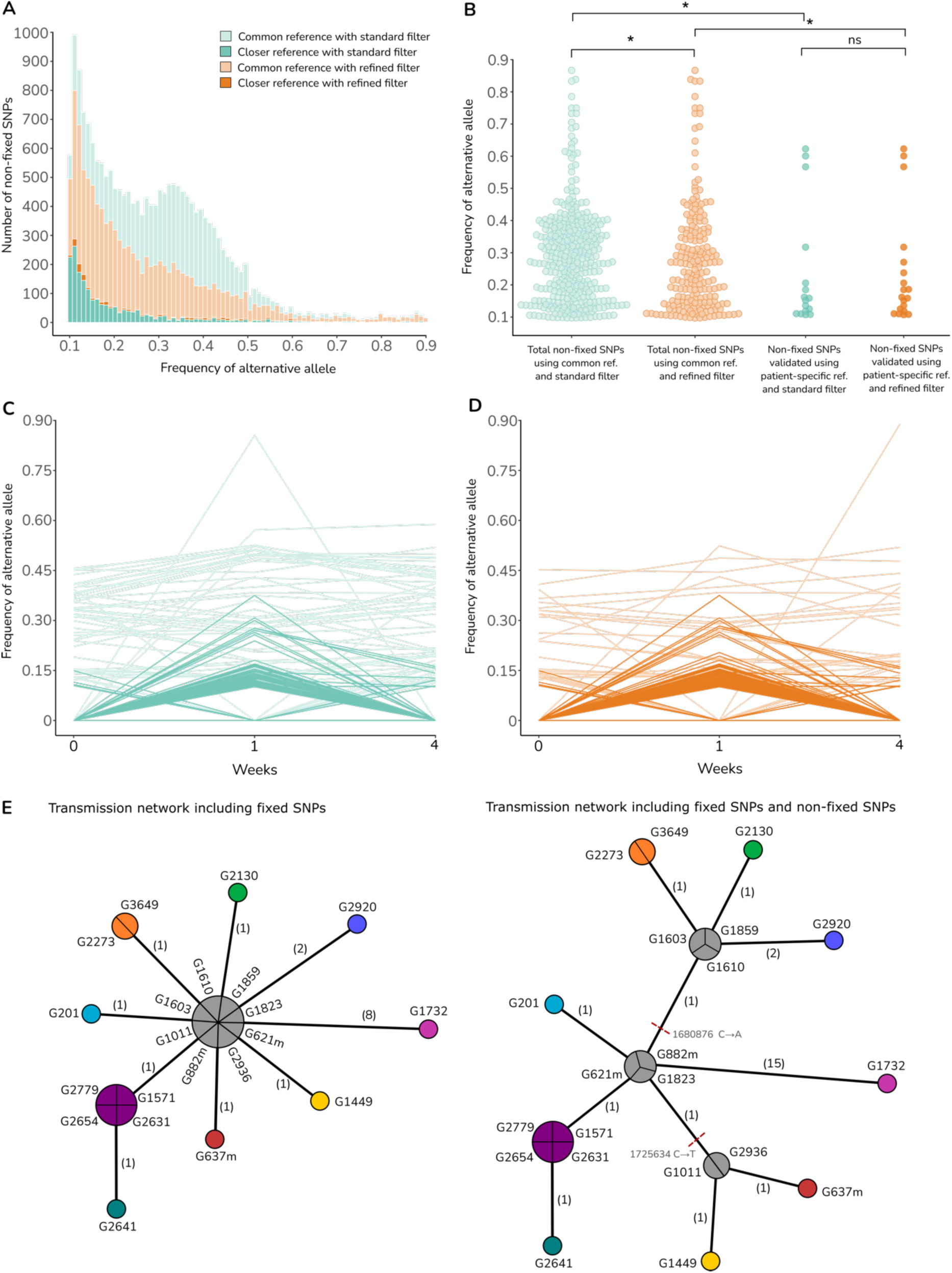
Differences between common and closer reference genome approaches. (A) Distributions of non-fixed SNPs detected in isolates from transmission clusters. Our standard short-read pipeline was performed twice in 372 MTB samples of 77 transmission clusters, using the MTBCA and a cluster-specific complete genome as reference genomes in each analysis. Additionally, standard and refined masking filters were applied in parallel to each reference approach. The distributions of non-fixed SNPs obtained by the common reference genome approach with standard and refined filters are plotted in pastel blue and pastel orange, respectively. The ones obtained by the cluster-specific reference with standard and refined filters are colored in blue and orange, respectively. (B) Distributions of non-fixed SNPs that were identified in four patients with two serial isolates. The figure shows on the left the distributions of total non-fixed SNPs detected by the common reference approach in pastel blue for the standard masking filter and in pastel orange for the refined one. On the right are displayed the distributions of validated non-fixed SNPs by the patient-specific reference approach for both types of filters (also see Table S2). ns, not significant; *, p < 0.05. (C, D) Trajectories of non-fixed SNPs from isolates of the same patient along three time points. The figures show the non-fixed SNPs detected by the common reference approach using the standard masking filter in blue (C) and the refined masking filter in orange (D). The frequencies (y-axis) are shown at 0, 1, and 4 weeks (x-axis). Only the non-fixed SNPs validated by the patient-specific reference approach are colored in intense blue or orange. It shows that most mutations at intermediate frequencies could not be validated. Notably, this was also observed with two reaching a frequency of 0.85 (also see Table S2). (E) Example of an original transmission network (left) and the same network including non-fixed SNPs (right). This figure illustrates how incorporating non-fixed SNPs (indicated by red dashed lines) into transmission analysis can enhance network resolution.

## DISCUSSION

In this study, we demonstrate that high-quality complete genomes provide a more comprehensive view of MTBC diversity, with important implications for both pathogen evolution and epidemiology. By combining our optimized DNA extraction protocol with HiFi sequencing, we can generate highly accurate long-read data to construct complete genomes without relying on short-read polishing. Complete genomes significantly increase the observed diversity at a global scale, uncovering previously overlooked variation that enhances our understanding of MTBC evolution, particularly the role of *pe/ppe* genes in host-pathogen interactions. While the increase in diversity is less pronounced at the epidemiological scale, it still improves the resolution of transmission networks. On the contrary, we observe a substantial downward revision of within-host diversity relative to previous estimates.

MTBC has been regarded as a monomorphic pathogen^83^. While this is largely still true, complete genomes depict considerably more variation than previous approaches, allowing the identification of diversity hotspots. Pairwise genome alignment comparisons reveal a median of 312 additional SNPs, with almost 90% of this variation occurring in the inaccessible regions. Consistently, the complete genome dataset yielded a substantially 1.5-fold higher estimated evolutionary rate, suggesting that complete genomes potentially refine molecular dating approaches. Our analysis provides additional sources of variation to explore the role of MTB genetic diversity in functional or clinical phenotypes.

As expected, most of the observed diversity was concentrated in several *pe/ppe* genes, whose high variability has been linked to potential roles in antigenic variation and immune evasion. However, these hypotheses are unconfirmed due to conflicting evidence and difficulties in reconstructing the evolutionary history of the entire *pe/ppe* repertoire^6,10,84,85^. Answering this longstanding question, our results reveal that PE/PPE epitopes follow the same evolutionary pattern previously observed in non-PE/PPE antigens, with epitopes hyperconserved in comparison with non-epitopes and full antigen sequences^5,7^. However, we observe gene conversion events introducing numerous amino acid changes in some epitopes of PPE18, PPE19, and PPE57 antigens, a key source of variation accurately captured by complete genomes. Remarkably, we found extensive gene conversion in *ppe18*, a subunit of the M72/AS01E vaccine antigen, which showed 49% protection in a phase IIb trial^86^. While most events occur outside epitope regions, some affect epitope sequences as well. Diversity in *ppe18* has been described before^13,85,87^, but only complete genomes have revealed the full extent of this diversity, making the *ppe18* region a diversity hotspot in MTBC (see Fig. 2, S2), and confirming that gene conversion, the leading mechanism of this variation, can also affect epitope regions. Our results highlight the need to evaluate the full diversity of antigens for vaccine development and the role of gene conversion in the evolution of these genes. Other structural variations may also have an impact on virulence, as illustrated by the extraordinary diversity found in the *ppe38-71* locus (Fig. 5).

At an epidemiological scale, our work establishes the first population-based benchmark for tracing genomic transmission by using complete genomes. Currently, genomic transmission links are defined by a 10-12 SNP threshold, and the low observed diversity precludes more detailed reconstruction of transmission networks^88,89^. Complete genome data offer a modest but significant gain in genetic distance between closely related samples, with a median of 1 additional SNP and 2 additional mutations in pairwise comparisons. Even more, this approach allows the confident incorporation of indel data in transmission analyses. Although indels have been traditionally excluded due to detection challenges, there is growing interest in developing more robust methods for detecting and integrating them, as they represent a potentially valuable source of information^90,91^. In our study, we reliably identified different types of indels, recovering genetic signals that are typically captured by classical genetic markers such as IS6110 mobile elements and variations in tandem repeats observed in spoligotyping and MIRU-VNTR. Occasionally, multiple SNPs can be gained from gene conversion events (as shown in Cluster 10)^13^. The inclusion of additional diversity affects the underlying genetic networks and alters the reconstruction of 20% of transmission histories (see Fig. S3). Additional research will be necessary to establish new SNP cutoffs and the integration of structural variants and other major sources of variation, such as gene conversion.

Complete genomes also impact the understanding of within-host variation, which is typically detected by identifying non-fixed variants within a bacterial population^82^. Although short-read data can accurately estimate variant frequencies, distinguishing genuine variants is challenging due to errors in mapping and calling to a common reference genome^92–94^. Here, we show that mapping short reads to a complete genome from the same patient vastly improves non-fixed variant detection removing up to 70-80% of variants from mapping to a common reference, depending on the masking filter. This strong filter effect highlights the relevance of applying refined filters^4,95^ when using a common reference. In contrast, minimal differences were observed when using a patient-specific reference, suggesting that masking filters may be unnecessary in this approach. These findings have profound implications for understanding the evolution of the pathogen during infection and treatment, and the emergence of drug resistance, calling into question previous estimates of within-host diversity. In addition, non-fixed variants may increase resolution in transmission networks^96–98^. In our population-based dataset, we could use them to increase the resolution in 23% transmission clusters, suggesting that cluster-specific references can pave the way for integrating non-fixed variants to clarify outbreaks.

Currently, long-read sequencing is not without caveats as it requires higher DNA quality and quantity than short-read sequencing, which is difficult to meet with mycobacteria. However, these requirements are expected to lessen shortly; meanwhile, we provide a fine-tuned protocol that has allowed us to recover complete genomes in almost all cases. Streamlining downstream analysis of complete genomes is also critically important; accurate multiple genome alignment remains a challenge^99^, especially for structural variants or high-complexity regions such as certain *pe*/*ppe* genes. We also acknowledge that the high enrichment of Lineage 4 in our dataset restricts the representation of global diversity. Even so, it still offers a detailed view of MTBC diversity across evolutionary scales and reveals the full potential of complete genomes. Future work with more diverse samples is needed to elucidate the role of MTBC diversity in host-pathogen interactions.

In summary, we present the first large-scale genomic study of *Mycobacterium tuberculosis* using complete genomes across evolutionary scales. Advances in long-read sequencing technologies suggest that *de novo* genome assembly will likely become routine in the future, and the resulting opportunities will present challenges in analysis and interpretation. Our work reveals substantial untapped genetic diversity that provides new avenues for understanding the ongoing evolution of *Mycobacterium tuberculosis* and how it is shaped by host-pathogen interactions. Furthermore, improving resolution at both the epidemiological and within-host levels will support the development of new strategies for TB control.

## DATA AVAILABILITY

All sequence data have been deposited on the European Nucleotide Archive in the projects PRJEB29604, PRJEB38719, PRJEB65844, PRJEB89456, PRJEB89397, PRJEB70424, and PRJEBXXXXX and are publicly available. The accession numbers for each sample are included in Tables S4, S5, and S12.

## CODE AVAILABILITY

All original code has been deposited at Zenodo with DOI:10.5281/zenodo.15489014 and is publicly available at https://github.com/anamatgu/MTBC_complete_genomes.

## CONSORTIA

The members of the Valencia Region TB Working Group are Aurora Blasco, Bárbara Gomila-Sard, José Luis López-Hontangas, Rafael Borrás, María Angeles Clari, Javier Colomina, David Navarro, Concepción Gimeno, María Remedio Guna-Serrano, Juan J Camarena, Ester Colomer-Roig, Nieves Orta, María M Ruiz-García, Nieves Gonzalo-Jiménez, Adelina Gimeno-Gascón, Juan Carlos Rodríguez, María Borrás-Máñez, Isabel Escribano, Olalla Martinez-Macias, and Oscar Esparcia-Rodríguez.

## ACKNOWLEDGMENTS

This project received funding from the European Research Council under the European Union’s Horizon 2020 Research and Innovation Programme (Grant 101001038, TB-RECONNECT) awarded to I.C. Additional support was provided by the Spanish Ministry of Science and Innovation (PID2022-137607OB-I00) and the Generalitat Valenciana (Project CIPROM/2023/30), both also awarded to IC. AGM has been supported by a ‘Formación de Profesorado Universitario’ (FPU) grant (FPU19/04562) from the Spanish Ministry of Universities, and the Health Research 2021 Programme (TB-TARGET_HR21-00415) by LaCaixa Foundation. Work in IC’s laboratory is further supported by the European Union’s Next Generation, through CSIC’s Global Health Platform (PTI Salud Global). JAR is hired under the Generation D initiative, promoted by Red.es, an organisation attached to the Ministry for Digital Transformation and the Civil Service, for the attraction and retention of talent through grants and training contracts (MMT24-PTI-SG-01), financed by the Recovery, Transformation, and Resilience Plan through the European Union’s Next Generation funds. We also gratefully acknowledge Jason Stout for his guidance during the manuscript revision.

## AUTHOR CONTRIBUTIONS

AMG conducted the formal analysis, curated, interpreted, and visualized the data. MMM, MH, ZI, MGL, and JAR assisted in the curation, formal analysis, and interpretation of the data. AMG and IC wrote the original draft. AMG, MTP, LMP, GDM, and MGL collected the data. AMG, MTP, and MGL validated the data. AGB, RB, MBM, JJC, JC, IE, OER, CG, AGG, BGS, NGJ, MRGS, JLLH, DN, MN, NO, JCR, MMRG, AB, MAG, MBA, and OMM provided resources. IC, FGC, and ZI assisted in the conceptualization of the study and supervised the process. IC acquired the funding and administered the project. All the authors contributed to the revision and editing of the paper, and all of them approved the final version and its publication.

## EXTENDED DATA

**Extended Data Table 1.**
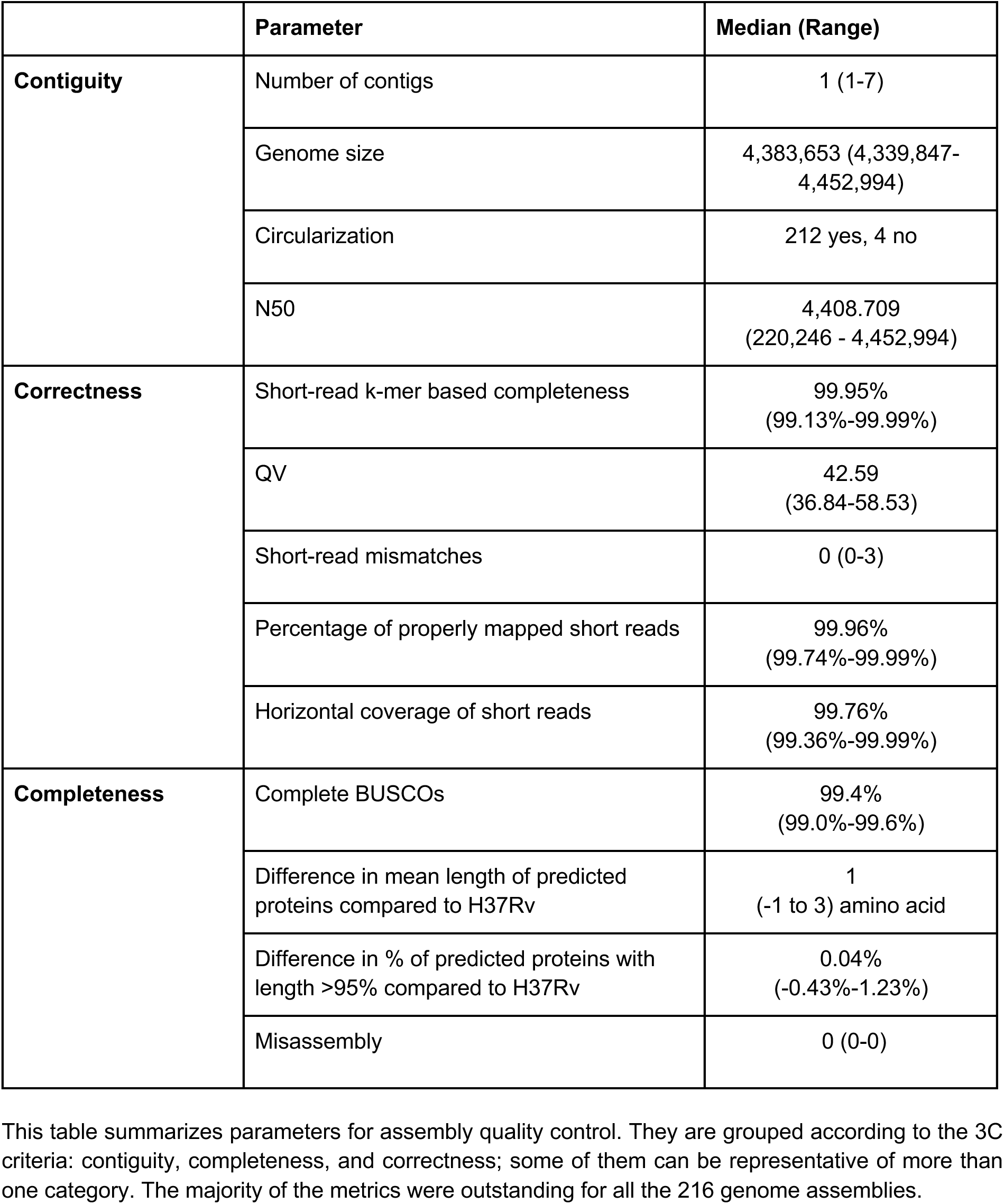
Quality control parameters of genome assemblies classified by the 3C criteria.

## REFERENCES

1. World Health Organization. Global Tuberculosis Report 2024. (World Health Organization, 2024).

2. Goig, G. A. et al. Ecology, global diversity and evolutionary mechanisms in the Mycobacterium tuberculosis complex. Nat Rev Microbiol (2025) doi:10.1038/s41579-025-01159-w.

3. Meehan, C. J. et al. Whole genome sequencing of Mycobacterium tuberculosis: current standards and open issues. Nat Rev Microbiol 17, 533–545 (2019).

4. Marin, M. et al. Benchmarking the empirical accuracy of short-read sequencing across the M. tuberculosis genome. Bioinformatics 38, 1781–1787 (2022).

5. Comas, I. et al. Human T cell epitopes of Mycobacterium tuberculosis are evolutionarily hyperconserved. Nat Genet 42, 498–503 (2010).

6. Copin, R. et al. Sequence Diversity in the pe_pgrs Genes of Mycobacterium tuberculosis Is Independent of Human T Cell Recognition. mBio (2014) doi:10.1128/mbio.00960-13.

7. Coscolla, M. et al. M. tuberculosis T Cell Epitope Analysis Reveals Paucity of Antigenic Variation and Identifies Rare Variable TB Antigens. Cell Host Microbe 18, 538–548 (2015).

8. Di Marco, F. et al. Advantages of long- and short-reads sequencing for the hybrid investigation of the genome. Front Microbiol 14, 1104456 (2023).

9. Bainomugisa, A. et al. A complete high-quality MinION nanopore assembly of an extensively drug-resistant Mycobacterium tuberculosis Beijing lineage strain identifies novel variation in repetitive PE/PPE gene regions. Microb Genom 4, (2018).

10. Phelan, J. E. et al. Recombination in pe/ppe genes contributes to genetic variation in Mycobacterium tuberculosis lineages. BMC Genomics 17, 151 (2016).

11. Gómez-González, P. J. et al. Functional genetic variation in / genes contributes to diversity in lineages and potential interactions with the human host. Front Microbiol 14, 1244319 (2023).

12. Hall, M. B. et al. Evaluation of Nanopore sequencing for Mycobacterium tuberculosis drug susceptibility testing and outbreak investigation: a genomic analysis. Lancet Microbe 4, e84–e92 (2023).

13. Stritt, C. et al. Gene conversion and duplication contribute to genetic variation in an outbreak of Mycobacterium tuberculosis. Microbial Genomics 11, 001396 (2025).

14. Method of the Year 2022: long-read sequencing. Nat Methods 20, 1 (2023).

15. Votintseva, A. A. et al. Same-Day Diagnostic and Surveillance Data for Tuberculosis via Whole-Genome Sequencing of Direct Respiratory Samples. J Clin Microbiol 55, 1285–1298 (2017).

16. Cancino-Muñoz, I. et al. Population-based sequencing of Mycobacterium tuberculosis reveals how current population dynamics are shaped by past epidemics. (2022) doi:10.7554/eLife.76605.

17. Fukasawa, Y., Ermini, L., Wang, H., Carty, K. & Cheung, M.-S. LongQC: A Quality Control Tool for Third Generation Sequencing Long Read Data. G3 (Bethesda) 10, 1193–1196 (2020).

18. Wood, D. E. & Salzberg, S. L. Kraken: ultrafast metagenomic sequence classification using exact alignments. Genome Biol 15, R46 (2014).

19. Lu, J. et al. Metagenome analysis using the Kraken software suite. Nat Protoc 17, 2815–2839 (2022).

20. Sim, S. B., Corpuz, R. L., Simmonds, T. J. & Geib, S. M. HiFiAdapterFilt, a memory efficient read processing pipeline, prevents occurrence of adapter sequence in PacBio HiFi reads and their negative impacts on genome assembly. BMC Genomics 23, 157 (2022).

21. Comas, I. Genome of the inferred most recent common ancestor of the Mycobacterium tuberculosis complex. doi:10.5281/zenodo.3497110.

22. Li, H. Minimap2: pairwise alignment for nucleotide sequences. Bioinformatics 34, 3094–3100 (2018).

23. GitHub - pysam-developers/pysam: Pysam is a Python package for reading, manipulating, and writing genomics data such as SAM/BAM/CRAM and VCF/BCF files. It’s a lightweight wrapper of the HTSlib API, the same one that powers samtools, bcftools, and tabix. *GitHub* https://github.com/pysam-developers/pysam.

24. García-Marín, A. M. anamatgu/MTBC_complete_genomes: MTBC_CG_v0.1. doi:10.5281/zenodo.15489014.

25. Kolmogorov, M., Yuan, J., Lin, Y. & Pevzner, P. A. Assembly of long, error-prone reads using repeat graphs. Nature Biotechnology 37, 540–546 (2019).

26. Hunt, M. et al. Circlator: automated circularization of genome assemblies using long sequencing reads. Genome Biol 16, 294 (2015).

27. GitHub - PacificBiosciences/pbmm2: A minimap2 frontend for PacBio native data formats. *GitHub* https://github.com/PacificBiosciences/pbmm2.

28. Wickham, H. ggplot2: Elegant Graphics for Data Analysis. (Springer Science & Business Media, 2009).

29. The R Project for Statistical Computing. https://www.R-project.org/.

30. Carver, T., Harris, S. R., Berriman, M., Parkhill, J. & McQuillan, J. A. Artemis: an integrated platform for visualization and analysis of high-throughput sequence-based experimental data. Bioinformatics 28, 464–469 (2012).

31. Garrison, E. & Marth, G. Haplotype-based variant detection from short-read sequencing. (2012).

32. Tan, A., Abecasis, G. R. & Kang, H. M. Unified representation of genetic variants. Bioinformatics 31, 2202–2204 (2015).

33. Shumate, A. & Salzberg, S. L. Liftoff: accurate mapping of gene annotations. Bioinformatics 37, 1639–1643 (2021).

34. Basic local alignment search tool. Journal of Molecular Biology 215, 403–410 (1990).

35. PacBio. Beyond contiguity — assessing the quality of genome assemblies with the 3 Cs. *PacBio* https://www.pacb.com/blog/beyond-contiguity/ (2020).

36. Molina-Mora, J. A., Campos-Sánchez, R., Rodríguez, C., Shi, L. & García, F. High quality 3C de novo assembly and annotation of a multidrug resistant ST-111 Pseudomonas aeruginosa genome: Benchmark of hybrid and non-hybrid assemblers. Scientific Reports 10, 1392 (2020).

37. Manni, M., Berkeley, M. R., Seppey, M., Simão, F. A. & Zdobnov, E. M. BUSCO Update: Novel and Streamlined Workflows along with Broader and Deeper Phylogenetic Coverage for Scoring of Eukaryotic, Prokaryotic, and Viral Genomes. Mol Biol Evol 38, 4647–4654 (2021).

38. GitHub - phiweger/ideel: Indels are not ideal - quick test for interrupted ORFs in bacterial/microbial genomes. GitHub https://github.com/phiweger/ideel.

39. Rhie, A., Walenz, B. P., Koren, S. & Phillippy, A. M. Merqury: reference-free quality, completeness, and phasing assessment for genome assemblies. Genome Biol 21, 245 (2020).

40. Hyatt, D. et al. Prodigal: prokaryotic gene recognition and translation initiation site identification. BMC Bioinformatics 11, 1–11 (2010).

41. Gurevich, A., Saveliev, V., Vyahhi, N. & Tesler, G. QUAST: quality assessment tool for genome assemblies. Bioinformatics 29, 1072–1075 (2013).

42. Smolka, M. et al. Detection of mosaic and population-level structural variants with Sniffles2. Nature Biotechnology 42, 1571–1580 (2024).

43. Hickey, G. et al. Pangenome graph construction from genome alignments with Minigraph-Cactus. Nature Biotechnology 42, 663–673 (2023).

44. Hickey, G., Paten, B., Earl, D., Zerbino, D. & Haussler, D. HAL: a hierarchical format for storing and analyzing multiple genome alignments. Bioinformatics 29, 1341–1342 (2013).

45. GitHub - mikolmogorov/maf2synteny: A tool for recovering synteny blocks from multiple alignment. GitHub https://github.com/mikolmogorov/maf2synteny.

46. Marçais, G. et al. MUMmer4: A fast and versatile genome alignment system. PLoS Comput Biol 14, e1005944 (2018).

47. Kurtz, S. et al. Versatile and open software for comparing large genomes. Genome Biol 5, R12 (2004).

48. TGU file repository. http://tgu.ibv.csic.es/?page_id=1794.

49. Li, H. & Durbin, R. Fast and accurate short read alignment with Burrows-Wheeler transform. Bioinformatics 25, 1754–1760 (2009).

50. Koboldt, D. C. et al. VarScan 2: somatic mutation and copy number alteration discovery in cancer by exome sequencing. Genome Res 22, 568–576 (2012).

51. Van der Auwera, G. A. & O’Connor, B. D. Genomics in the Cloud: Using Docker, GATK, and WDL in Terra. (O’Reilly Media, 2020).

52. Cingolani, P. et al. A program for annotating and predicting the effects of single nucleotide polymorphisms, SnpEff. Fly (2012) doi:10.4161/fly.19695.

53. Catalogue of mutations in Mycobacterium tuberculosis complex and their association with drug resistance. https://www.who.int/publications/i/item/9789240028173 (2021).

54. Page, A. J. et al. Snp-sites: rapid efficient extraction of SNPs from multi-FASTA alignments. Microb Genom 2, e000056 (2016).

55. Leigh, J. W. & Bryant, D. popart: full-feature software for haplotype network construction. Methods in Ecology and Evolution 6, 1110–1116 (2015).

56. Paradis, E. & Schliep, K. ape 5.0: an environment for modern phylogenetics and evolutionary analyses in R. Bioinformatics 35, 526–528 (2019).

57. Minh, B. Q. et al. IQ-TREE 2: New Models and Efficient Methods for Phylogenetic Inference in the Genomic Era. Mol Biol Evol 37, 1530–1534 (2020).

58. Sagulenko, P., Puller, V. & Neher, R. A. TreeTime: Maximum-likelihood phylodynamic analysis. Virus Evol 4, vex042 (2018).

59. Bouckaert, R. et al. BEAST 2.5: An advanced software platform for Bayesian evolutionary analysis. PLOS Computational Biology 15, e1006650 (2019).

60. Bos, K. I. et al. Pre-Columbian mycobacterial genomes reveal seals as a source of New World human tuberculosis. Nature 514, (2014).

61. Sabin, S. et al. A seventeenth-century Mycobacterium tuberculosis genome supports a Neolithic emergence of the Mycobacterium tuberculosis complex. Genome Biology 21, 1–24 (2020).

62. Correcting for constant sites in BEAST2. https://groups.google.com/g/beast-users/c/QfBHMOqImFE.

63. Rambaut, A., Drummond, A. J., Xie, D., Baele, G. & Suchard, M. A. Posterior Summarization in Bayesian Phylogenetics Using Tracer 1.7. Syst Biol 67, 901–904 (2018).

64. Wickham, H. et al. Welcome to the Tidyverse. Journal of Open Source Software 4, 1686 (2019).

65. GitHub - wilkelab/ggridges: Ridgeline plots in ggplot2. *GitHub* https://github.com/wilkelab/ggridges.

66. Vilella, A. J., Blanco-Garcia, A., Hutter, S. & Rozas, J. VariScan: Analysis of evolutionary patterns from large-scale DNA sequence polymorphism data. Bioinformatics 21, 2791–2793 (2005).

67. Cortes, T. et al. Genome-wide mapping of transcriptional start sites defines an extensive leaderless transcriptome in Mycobacterium tuberculosis. Cell Rep 5, 1121–1131 (2013).

68. Ranwez, V., Harispe, S., Delsuc, F. & Douzery, E. J. P. MACSE: Multiple Alignment of Coding SEquences accounting for frameshifts and stop codons. PLoS One 6, e22594 (2011).

69. Ranwez, V., Douzery, E. J. P., Cambon, C., Chantret, N. & Delsuc, F. MACSE v2: Toolkit for the Alignment of Coding Sequences Accounting for Frameshifts and Stop Codons. Mol Biol Evol 35, 2582–2584 (2018).

70. Croucher, N. J. et al. Rapid phylogenetic analysis of large samples of recombinant bacterial whole genome sequences using Gubbins. Nucleic Acids Res 43, e15 (2015).

71. GitHub - Pas-Kapli/FastaCon: concatenates multiple fasta files in a new fasta file. *GitHub* https://github.com/Pas-Kapli/FastaCon.

72. Ates, L. S. New insights into the mycobacterial PE and PPE proteins provide a framework for future research. Mol Microbiol 113, 4–21 (2020).

73. IEDB.org: Free epitope database and prediction resource. https://www.iedb.org/.

74. Rodrigo, A. G. & Learn, G. H., Jr. Computational and Evolutionary Analysis of HIV Molecular Sequences. (Springer Science & Business Media, 2007).

75. Nei, M. & Gojobori, T. Simple methods for estimating the numbers of synonymous and nonsynonymous nucleotide substitutions. Mol Biol Evol 3, 418–426 (1986).

76. Didelot, X. & Parkhill, J. A scalable analytical approach from bacterial genomes to epidemiology. Philosophical Transactions of the Royal Society B (2022) doi:10.1098/rstb.2021.0246.

77. Hunt, M. et al. Minos: variant adjudication and joint genotyping of cohorts of bacterial genomes. Genome Biology 23, 1–23 (2022).

78. Darling, A. E., Mau, B. & Perna, N. T. progressiveMauve: multiple genome alignment with gene gain, loss and rearrangement. PLoS One 5, e11147 (2010).

79. Lindenbaum, P. JVarkit: java-based utilities for Bioinformatics. (2015) doi:10.6084/m9.figshare.1425030.v1.

80. Khelik, K., Lagesen, K., Sandve, G. K., Rognes, T. & Nederbragt, A. J. NucDiff: in-depth characterization and annotation of differences between two sets of DNA sequences. BMC Bioinformatics 18, 1–14 (2017).

81. Mariner-Llicer, C. et al. Genetic diversity within diagnostic sputum samples is mirrored in the culture of Mycobacterium tuberculosis across different settings. Nat Commun 15, 7114 (2024).

82. Martin, M. A., Lee, R. S., Cowley, L. A., Gardy, J. L. & Hanage, W. P. Within-host Mycobacterium tuberculosis diversity and its utility for inferences of transmission. Microbial Genomics 4, e000217 (2018).

83. Stritt, C. & Gagneux, S. How do monomorphic bacteria evolve? The Mycobacterium tuberculosis complex and the awkward population genetics of extreme clonality. Peer Community J. 3, (2023).

84. Fishbein, S., van Wyk, N., Warren, R. M. & Sampson, S. L. Phylogeny to function: PE/PPE protein evolution and impact on Mycobacterium tuberculosis pathogenicity. Mol Microbiol 96, 901–916 (2015).

85. McEvoy, C. R. E. et al. Comparative Analysis of Mycobacterium tuberculosis pe and ppe Genes Reveals High Sequence Variation and an Apparent Absence of Selective Constraints. PLOS ONE 7, e30593 (2012).

86. Dale, K. D. & Denholm, J. T. Optimising the M72/AS01E tuberculosis vaccine candidate phase 3 trial based on the phase 2b trial results. Vaccine 49, 126816 (2025).

87. Homolka, S., Ubben, T. & Niemann, S. High Sequence Variability of the ppE18 Gene of Clinical Mycobacterium tuberculosis Complex Strains Potentially Impacts Effectivity of Vaccine Candidate M72/AS01E. PLoS One 11, e0152200 (2016).

88. Walker, T. M. et al. Assessment of Mycobacterium tuberculosis transmission in Oxfordshire, UK, 2007-12, with whole pathogen genome sequences: an observational study. Lancet Respir Med 2, 285–292 (2014).

89. Xu, Y. et al. High-resolution mapping of tuberculosis transmission: Whole genome sequencing and phylogenetic modelling of a cohort from Valencia Region, Spain. PLOS Medicine 16, e1002961 (2019).

90. Worakitchanon, W. et al. Comprehensive analysis of Mycobacterium tuberculosis genomes reveals genetic variations in bacterial virulence. Cell host & microbe 32, (2024).

91. Dixit, A. et al. Whole genome sequencing identifies bacterial factors affecting transmission of multidrug-resistant tuberculosis in a high-prevalence setting. Scientific Reports 9, 1–10 (2019).

92. Lee, R. S. & Behr, M. A. Does Choice Matter? Reference-Based Alignment for Molecular Epidemiology of Tuberculosis. Journal of Clinical Microbiology (2016) doi:10.1128/jcm.00364-16.

93. Bush, S. J. et al. Genomic diversity affects the accuracy of bacterial single-nucleotide polymorphism-calling pipelines. Gigascience 9, (2020).

94. Valiente-Mullor, C. et al. One is not enough: On the effects of reference genome for the mapping and subsequent analyses of short-reads. PLOS Computational Biology 17, e1008678 (2021).

95. Modlin, S. J. et al. Exact mapping of Illumina blind spots in the Mycobacterium tuberculosis genome reveals platform-wide and workflow-specific biases. Microbial Genomics 7, 000465 (2021).

96. López, M. G. et al. Deciphering the Tangible Spatio-Temporal Spread of a 25-Year Tuberculosis Outbreak Boosted by Social Determinants. Microbiology Spectrum (2023) doi:10.1128/spectrum.02826-22.

97. Pérez-Llanos, F. J. et al. Transmission Dynamics of a Mycobacterium tuberculosis Complex Outbreak in an Indigenous Population in the Colombian Amazon Region. Microbiology spectrum 11, (2023).

98. Lee, R. S., Proulx, J.-F., McIntosh, F., Behr, M. A. & Hanage, W. P. Previously undetected super-spreading of Mycobacterium tuberculosis revealed by deep sequencing. (2020) doi:10.7554/eLife.53245.

99. Armstrong, J. et al. Progressive Cactus is a multiple-genome aligner for the thousand-genome era. Nature 587, 246–251 (2020).

