## Supplementary_Information for "Complete genomes reveal the full extent of *Mycobacterium tuberculosis* complex diversity across evolutionary scales"

**Supplementary Notes**

- High-quality MTBC complete genomes obtained by long-read HiFi sequencing
- Basic statistics of the Multiple Genome Alignment
- Fixed SNPs missed by short-read mapping and assessment of masked regions
- Mapping to a closer reference genome enhances the quality of short-read alignment

**Supplementary Figures**

- Figure S1. Basic statistics of long-read sequencing and features of the dataset
- Figure S2. Intragenic nucleotide diversity of six highly variable *pe/ppe* genes in *M. tuberculosis*
- Figure S3. Transmission network diagrams of all transmission clusters

**Supplementary Tables**

- Table S1. Functional enrichment analysis of genes with diversity hotspots
- Table S2. Number of non-fixed SNPs detected in intrapatient samples using the MTBCA reference genome and patient-specific reference genomes

#### Supplementary Notes

##### High-quality MTBC complete genomes obtained by long-read HiFi sequencing

Of 266 MTBC culture-positive isolates available in the Valencia Region of Spain from 2016, 216 (80%) were sequenced using the PacBio HiFi method. DNA from 162 isolates (75%) was sequenced from the archive aliquots, while 54 isolates (25%) required regrowth to obtain DNA suitable for long-read sequencing. An average sequencing depth of 100.5x was achieved with an average of 98,930 reads per isolate, a mean read length of 4,505, and a rate of 0.0005 single-mismatches per base (Fig. S1, Table S3).

We obtained 216 draft genomes with a median coverage of 90 (22–353). Due to inconsistencies in circularization with Flye, we closed draft assemblies with Circlator, resulting in changes to 49 assemblies. Collapsed repeats were fixed in 6 assemblies with manual curation. In the self-correction polishing step, 45 changes were introduced across 28 assemblies, including 29 one- or two-base pair deletions, 11 one-base pair insertions, and 5 SNPs.

We achieved 212 genome assemblies closed in a single circular contig with a median length/N50 of 4,408,982 (4,339,847–4,452,994) base pairs. Additionally, all 216 genome assemblies were highly accurate and complete (Table S1, Table S5). In terms of concordance with short-read data, all 216 assemblies had: i) over 99.9% k-mer based completeness, ii) a QV range of Q37 to Q59 (indicating an accuracy above 99.9%), and iii) a median of 0 (0–3) mismatches, over 99.7% mapped short reads, and a horizontal coverage greater than 99.4%.

The assembly and long-reads metrics were also outstanding: i) 99% of complete BUSCOs were found in all genome assemblies, with only 0.4% or fewer being fragmented and less than 0.7% missing, ii) the mean length of proteins predicted by Prodigal differed by a median of 1 (-1 to 3) amino acid compared to the H37Rv reference genome, and the proportion of proteins predicted by IDEEL with lengths above 95% differed by only 0.06% on average compared to the H37Rv reference genome, and iii) only non-fixed structural variations were found, indicating no misassemblies according to long-read data. All parameters used to assess contiguity, completeness, and correctness can be found in Table S5. As all 216 assemblies demonstrated consistently high-quality metrics, they were all included in downstream analyses.

##### Basic statistics of the Multiple Genome Alignment

We generated a Multiple Genome Alignment (MGA) encompassing all 216 Mtbc genome assemblies. The MGA consisted of 1519 blocks with an average block degree of 210 sequences. From this whole genome alignment, 35 synteny blocks were identified, with an average block degree of 214 sequences and a mean horizontal coverage of 90.7% per sequence (median of 90.9%).

During the refinement of the MGA, we identified 28710 non-redundant positions that harbored a SNP in at least one sample. Of those, 25987 positions were validated. Of the 2154 non-validated positions, 2154 had calls only detected by cactus and thus only masked in the specific samples that harbored them, and 569 were considered ambiguous positions and masked in all the samples as they had a variable number of calls depending on the approach. Additionally, 122 positions with non-fixed variants were masked in the specific samples.

##### Fixed SNPs missed by short-read mapping and assessment of masked regions

When comparing pairwise genetic distances from short-read and complete genome data, there were 5 fixed SNPs missed by the short-read approach in preserved regions for different causes: i) three

SNPs were missed due to mapping errors in regions with structural variants (duplications), ii) one SNP was located in an insertion within an intergenic region shared by both complete genomes but absent in the MTBCA reference genome, iii) two SNPs were detected but did not meet the required coverage threshold, and iv) one SNP was detected but did not reach the allele frequency threshold (AF > 0.9). As expected, indels were almost evenly distributed, with 41% in masked regions and 59% in unmasked regions (Figure 5c).

Given the potential of variants in masked regions to enhance epidemiological resolution, we assessed the reliability of variant calling in these regions with the short-read mapping approach. We analyzed 1,081 polymorphic positions across 145 genes and 51 intergenic regions, with a detailed list provided in Table S10. The F1 score distribution was bimodal, with 50.3% (544/1,081) of positions exhibiting an F1 score of 0 and 12.8% (138/1,081) an F1 score of 1. Notably, positions with an F1 score of 0 were concentrated within 60 of the 161 analyzed regions, whereas positions with an F1 score of 1 were more widely distributed across 101 regions. However, both values coexisted in 13 regions, suggesting the need for annotation filters to selectively mask individual positions rather than entire regions.

**Mapping to a closer reference genome enhances the quality of short-read alignment**

We observe that mapping to a closer reference genome increases the reliability of mapping and thus subsequent calling steps compared to a common reference genome. This was supported by the following mapping quality statistics: (i) higher horizontal coverage, with a median difference of 0.6% (98.12% vs.97.53%), (ii) higher proportion of total mapped reads, with a median difference of 0.3% (97.83% vs. 97.53%), (iii) lower proportion of supplementary alignments, with a median difference of 0.06% (0.80% vs. 0.86%), and (iv) lower proportion of improperly paired mapped reads, with a median difference of 0.08% (0.58% vs. 0.66%). All differences were statistically significant ($p < 0.01$, Wilcoxon signed-rank test).

**Supplementary figures**

**Figure S1. Basic statistics of long-read sequencing and features of the dataset.**

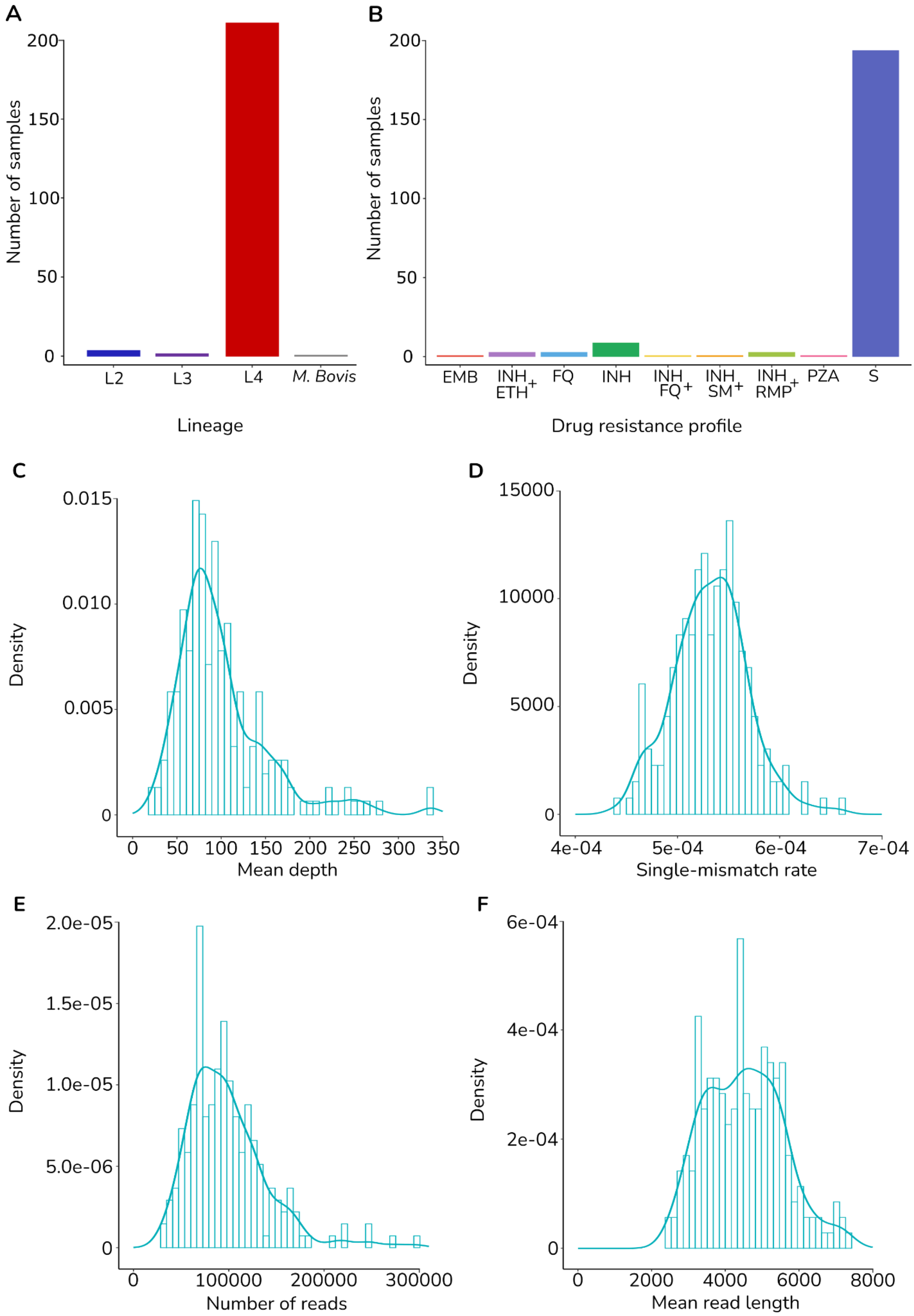


This figure shows the features of the dataset and the basic statistics used as quality control of the 216 isolates sequenced by HiFi technology:

(A) Frequency of lineages: L2 2.0% (4), L3 0.9% (2), L4 96.7% (209), and *M. bovis* 0.4% (1)

(B) Frequency of drug-resistance profiles: 89.8% (194) were pansusceptible, 6.5% (14) exhibited mono-resistance (INH: 9; EMB: 1; PZA: 1; FQ: 3), and 3.7% (8) showed resistance to two drugs (INH+RMP: 3; EMB+INH: 3; INH+FQ: 1; INH+SM: 1) of the 216 MTBC isolates with matched short-read and complete genome data.
(C) Distribution of mean depth achieved in each isolate, with an average depth of 100.5x
(D) Distribution of the single-mismatch rate per isolate, with an average of 0.00053
(E) Distribution of the total number of reads reached per isolate, with an average of 98,930 reads
(F) Distribution of the mean read length per isolate, with an average of 4,505bp.

INH= Isoniazid; RMP = Rifampicin; EBM = Ethambutol; PZA = Pyrazinamide; SM = Streptomycin; FQ = Fluoroquinolones; ETH = Ethionamide

**Figure S2. Intragenic nucleotide diversity of six highly variable *pe/ppe* genes in *M. tuberculosis.***

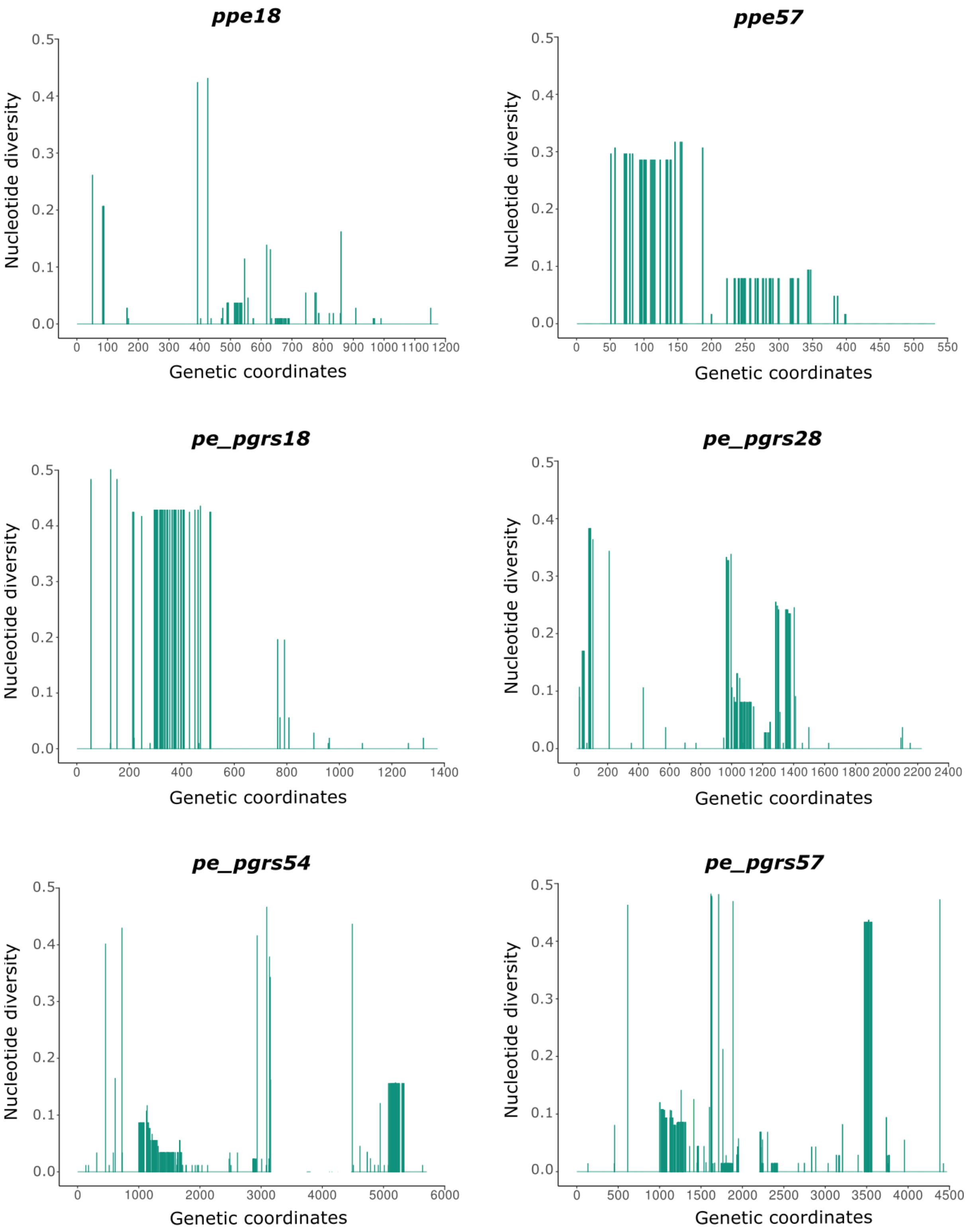


Six of the most genetically diverse *pe/ppe* genes were selected to illustrate patterns of intragenic diversity. Each panel displays nucleotide diversity (x-axis) along the full length of the gene (y-axis). Despite overall high genetic diversity, variation is not equally distributed, with specific regions within each gene showing higher rates of nucleotide diversity.

### Figure S3. Transmission network diagrams of 24 clusters identified using a 10-SNP threshold

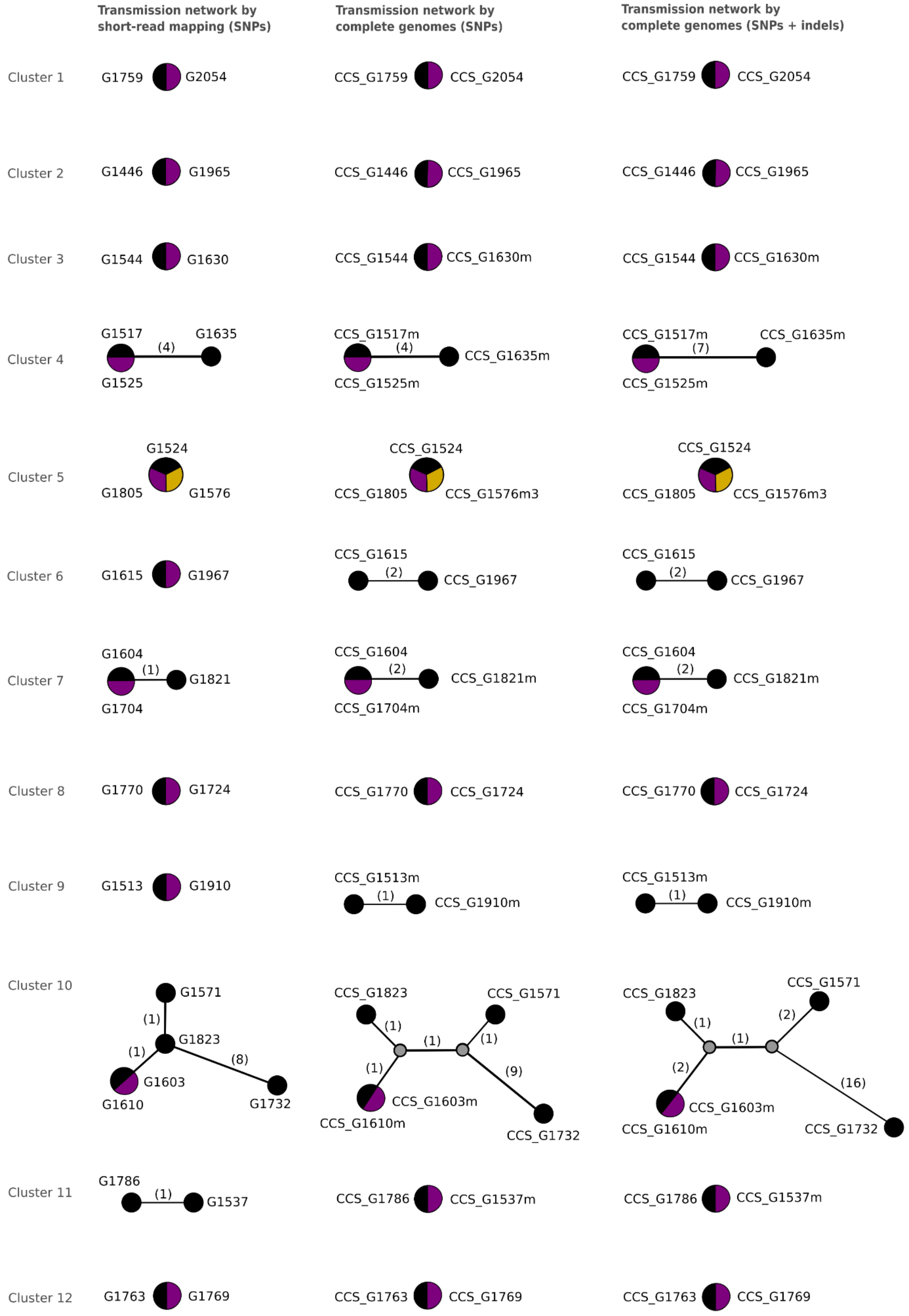

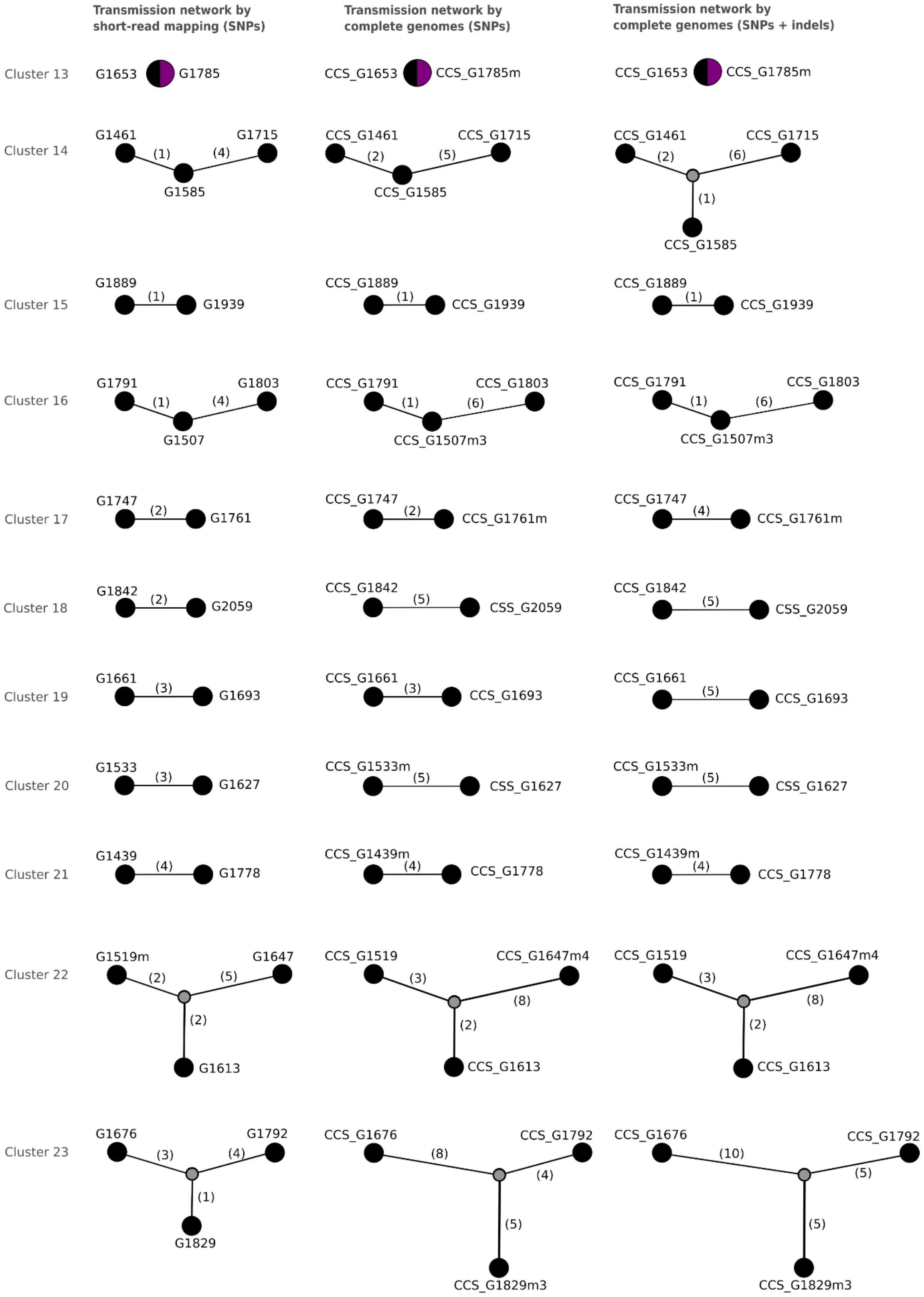
Transmission network by short-read mapping (SNPs)
Transmission network by complete genomes (SNPs)
Transmission network by complete genomes (SNPs + indels)
Cluster 13
G1653
G1785
CCS_G1653
CCS_G1785m
CCS_G1653
CCS_G1785m
Cluster 14
G1461
(1)
(4)
G1715
G1585
CCS_G1461
(2)
(5)
CCS_G1715
CCS_G1585
CCS_G1461
(2)
(6)
CCS_G1715
(1)
CCS_G1585
Cluster 15
G1889
(1)
G1939
CCS_G1889
(1)
CCS_G1939
CCS_G1889
(1)
CCS_G1939
Cluster 16
G1791
(1)
(4)
G1803
G1507
CCS_G1791
(1)
(6)
CCS_G1803
CCS_G1507m3
CCS_G1791
(1)
(6)
CCS_G1803
CCS_G1507m3
Cluster 17
G1747
(2)
G1761
CCS_G1747
(2)
CCS_G1761m
CCS_G1747
(4)
CCS_G1761m
Cluster 18
G1842
(2)
G2059
CCS_G1842
(5)
CSS_G2059
CCS_G1842
(5)
CSS_G2059
Cluster 19
G1661
(3)
G1693
CCS_G1661
(3)
CCS_G1693
CCS_G1661
(5)
CCS_G1693
Cluster 20
G1533
(3)
G1627
CCS_G1533m
(5)
CSS_G1627
CCS_G1533m
(5)
CSS_G1627
Cluster 21
G1439
(4)
G1778
CCS_G1439m
(4)
CCS_G1778
CCS_G1439m
(4)
CCS_G1778
Cluster 22
G1519m
(2)
(5)
G1647
(2)
G1613
CCS_G1519
(3)
CCS_G1647m4
(8)
(2)
CCS_G1613
CCS_G1519
(3)
CCS_G1647m4
(8)
(2)
CCS_G1613
Cluster 23
G1676
(3)
(4)
G1792
(1)
G1829
CCS_G1676
(8)
CCS_G1792
(4)
(5)
CCS_G1829m3
CCS_G1676
(10)
CCS_G1792
(5)
(5)
CCS_G1829m3
Cluster 24
G1835
(9)
G1851
CCS_G1835
(11)
CCS_G1851
CCS_G1835
(11)
CCS_G1851

Twenty-four transmission clusters were identified in the 2016 dataset using standard short-read mapping analysis. The figure presents transmission networks for all clusters, comparing three data types: (i) SNPs from short-read data (first column), (ii) SNPs from complete genomes (second column), and (iii) SNPs and InDels from complete genomes (third column). Black bubbles represent external nodes (sampled isolates), grey bubbles indicate hypothetical nodes (unsampled intermediates), and samples with 0 SNP distance are grouped in larger bubbles colored purple or yellow to distinguish different isolates.

**Supplementary tables**

**Table S1. Functional enrichment analysis of genes with diversity hotspots.**

| COG id | COG category | Number of genes | Frequency | p-value | adjusted p-value |
|---|---|---|---|---|---|
| I.B | Energy Metabolism | 2 | 3.1 | 0.327 | 0.409 |
| I.I | Polyketide and non-ribosomal peptide synthesis | 2 | 3.1 | 0.150 | 0.245 |
| I.J | Broad regulatory functions | 3 | 4.7 | 1.000 | 1.000 |
| II.C | Cell envelope | 2 | 3.1 | 0.172 | 0.245 |
| IV.B | IS elements, Repeated sequences, and Phage | 11 | 17.2 | 1.37E-05 | 6.86E-05 |
| IV.C | PE and PPE families | 23 | 35.9 | 2.58E-15 | 2.58E-14 |
| IV.J | Cyclases | 2 | 3.1 | 0.007 | 0.022 |
| V | Conserved hypotheticals | 11 | 17.2 | 0.538 | 0.597 |
| VI | Unknowns | 3 | 4.7 | 0.020 | 0.050 |
| VII | Toxin Antitoxin System | 5 | 7.8 | 0.050 | 0.099 |

COG = Clusters of Orthologous Groups

**Table S2. Number of non-fixed SNPs detected in serial isolates using the MTBCA and patient-specific reference genomes with standard and refined masking filters.**

| | | Number of non-fixed SNPs | | | | | | | |
|---|---|---|---|---|---|---|---|---|---|
| Patient | Week | MTBCA reference + standard filter | Patient reference + standard filter | Validated by both references + standard filter | True positive nfSNPs by standard filter | MTBCA referenc e + refined filter | Patient reference + refined filter | Validated by both references + refined filter | True positive nfSNPs by refined filter |
| 1.0 | 0 | 33 | 1 | 1 | 3.0% | 14 | 1 | 1 | 7.1% |
| 1.1 | 4 | 30 | 0 | 0 | 0.0% | 11 | 0 | 0 | 0.0% |
| 2.0 | 0 | 52 | 4 | 4 | 7.7% | 37 | 4 | 4 | 10.8% |
| 2.1 | 1 | 140 | 89 | 88 | 62.9% | 113 | 90 | 88 | 77.9% |
| 2.2 | 4 | 69 | 9 | 8 | 11.6% | 38 | 10 | 9 | 23.7% |
| 3.0 | 0 | 86 | 1 | 1 | 1.2% | 32 | 4 | 3 | 9.4% |
| 3.1 | 4 | 65 | 3 | 3 | 4.6% | 57 | 3 | 1 | 1.8% |
| 4.0 | 0 | 39 | 1 | 1 | 2.6% | 25 | 6 | 6 | 24.0% |
| 4.1 | 2 | 47 | 5 | 5 | 10.6% | 28 | 1 | 1 | 3.6% |
| 5.0 | 0 | 40 | 3 | 3 | 7.5% | 17 | 2 | 2 | 11.8% |
| 5.1 | 6 | 39 | 2 | 2 | 5.1% | 24 | 5 | 5 | 20.8% |

nfSNP = non-fixed SNP; MTBCA = Mycobacterium tuberculosis complex most likely common ancestor
